# Transcriptome analysis of protein kinase MoCK2 affect acetyl-CoA metabolism and import of CK2 interacting mitochondrial proteins into mitochondria in rice blast fungus *Magnaporthe oryzae*

**DOI:** 10.1101/2022.07.28.501933

**Authors:** Lianhu Zhang, Chonglei Shan, Yifan Zhang, Wenjing Miao, Xiaoli Bing, Weigang Kuang, Zonghua Wang, Ruqiang Cui, Stefan Olsson

**Author notes:** These authors contributed equally (Lianhu Zhang, Chonglei Shan). Corresponding authors. E-mails: Ruqiang Cui,; Stefan Olsson.

## Abstract

The rice pathogen *Magnaporthe oryzae* causes severe losses to rice production. Previous studies have shown protein kinase MoCK2 essential for pathogenesis, and this ubiquitous eukaryotic might affect several processes in the fungus needed for infection. To better understand which cellular processes are affected by the MoCK2 activity, we performed a detailed RNAseq analysis of deletions MoCK2-beta1 and beta2 components in relation to the background strain Ku80 and connected this analysis with the abundance of substrates for proteins in a previous pulldown of the essential CKa subunit of CK2 to estimate effects on proteins directly interacting with CK2. The results showed that MoCK2 seriously affected carbohydrate metabolism, fatty acid metabolism, amino acid metabolism and related transporters and reduced acetyl-CoA production. CK2 phosphorylation can affect the folding of proteins and especially the effective formation of protein complexes by intrinsically disordered or mitochondrial import by destabilizing soluble alpha helixes. Upregulated genes found in the pulldown of the b1 and b2 mutants indicate that proteins directly interacting with CK2 are compensatory upregulated depending on their pulldown. A similar correlation was found for mitochondrial proteins. Taken together, the classes of proteins and the change in regulation in the b1 and b2 mutants suggest that CK2 has a central role in mitochondrial metabolism, secondary metabolism, and ROS resistance, in addition to the previously suggested role in the formation of new ribosomes, all processes central to efficient non-self responses as innate immunity.

**Importance:** The protein kinase CK2 is highly expressed and essential for plants, animals, and fungi affecting fatty acid-related metabolism. In addition, it directly affects the import of essential mitochondrial proteins into mitochondria. These effects mean CK2 is essential for lipid metabolism and mitochondrial function and, as shown before, crucial for making new translation machinery proteins. Taken together, our new results combined with previous published indicate that CK2 is an essential protein necessary for the capacity to launch efficient innate immunity responses and withstand the negative effect of such responses necessary for general resistance against invading bacteria and viruses as well as to interact with plants and withstand plant immunity responses and kill plant cells.

Protein kinase CK2, highly expressed and essential for plants, animals, and fungi, affects fatty acid-related metabolism and mitochondrial proteins, making it essential for the capacity to launch efficient innate immunity responses and plant pathogenicity

## Introduction

Rice is the most important food crop in the world. Most people in the world have rice as one of their staple foods. Thus rice production plays a vital role in world food security(1). However, huge losses are caused by diseases and pests every year. Among these diseases, rice blast is the most important, seriously affecting rice yield and quality (2), making prevention, damage limitation and controlling of rice blast infections essential to secure food supply. Presently, the use of resistant plant varieties is the most common method to prevent and control the occurrence of this disease (3). However, with the common practice of crop monoculture of plant cultivars and the dynamic genetic and epigenetic changes in physiological races of pathogens, the loss of resistance against pathogens of plant cultivars is a constant problem and threat (4). Therefore, increased basic knowledge of the mechanism of pathogenicity of the rice blast fungus is another important road towards the goal of preventing, limiting damage and hindering disease epidemics with devastating consequences.

The rice blast disease is caused by the fungus *Magnaporthe oryzae* (5). The infection develops as follows; conidia are spread by wind, attach on rice leaves, and germinate to form a germ tube. The tip of the germ tube expands to form an appressorium(6). Deposition of melanin on the cell wall makes it strong and, together with the accumulation of a large amount of glycerol in the appressorium, a huge turgor develops(7). The maturation of the appressorium is accompanied by autophagy of the conidial cells and nuclear mitotic divisions in the germ tube(8, 9). A penetration peg is formed at the bottom of mature appressorium that pierces the rice epidermal cells to complete the infection and enter the plant epidermal cell interior(10). Under suitable local conditions for pathogen invasion, rice leaves form disease spots after 5-7 days and then produce new conidia. Conidia spread and start a new disease cycle if spread to uninfected plants by the power of wind and rain(11). Previous research has shown that the infection process of *M. oryzae* is regulated by a series of signals, such as through the cAMP pathway and MAPK pathway, affecting the production and germination of conidia and the formation and infection by appressoria(12, 13). Deletion mutants of some important genes, such as PKA(12), (13) and the septin proteins involved in septal pore formation and dynamics (14), seriously hindered the infection and pathogenicity of *M. oryzae.* In our lab research, we cloned the protein kinase MoCK2. The protein kinase MoCK2 holoenzyme is an enzyme complex containing two catalytic subunits and two regulatory subunits. The catalytic subunit is encoded by one gene, MGG_03696, named, MoCKa. Two different genes encode the two regulatory subunits, MGG_ 00446 and MGG_ 05651, named MoCKb1 and MoCKb2, respectively. MoCKa deletions is lethal, and mutants cannot be obtained. The deletion mutants of the two regulatory subunits MoCKb1 and MoCKb2 showed severe phenotypic defects *in vitro* (we now use b1 as short for the deletion mutant of MoCKb1 and b2 for the deletion mutant of MoCKb2.). Both mutants b1 and b2 grew slowly, decreased spore production severely and lost pathogenicity. That was accompanied by a complete loss of capacity for infection for rice leaves, indicating that the MoCK2 holoenzyme supplies the pathogen with important functions needed for its pathogenic life-cycle (15). We further screened thousands of interacting proteins of MoCKa in a pulldown experiment. That analysis showed that the phosphorylation of MoCK2 and the accompanying dephosphorylation have important functions temporarily destabilizing alpha helixes, allowing the normal formation of IDP (intrinsically disordered proteins) protein-protein and protein polymer interactions, like the efficient formation of new ribosomes (16).

Based on this, we aimed to more accurately understand how the deletions of the MoCK2 components affect the activities of *M. oryzae* through transcriptome analysis of the b1 and b2 compared with the background strains Ku80 under *in vitro* conditions. An RNA-seq employing second-generation sequencing was performed to analyze the transcriptome differences between the mutants b1, b2, and the background strains Ku80 to get an idea of which processes are affected in the mutant and why we observed particular phenotypes

Transcriptome sequencing technology has been widely used in scientific research, including in the interaction between rice and *M. oryzae.* Gene expression changes in susceptible and resistant rice varieties infected by *M. oryzae* are analyzed by transcriptome sequencing and play important guiding roles in developing new rice varieties(17).

Currently, RNAseq is an often used convenient means to get indications of extended roles of genes and a comprehensive understanding of genes in the whole genome affected by a target gene’s direct and indirect activities. It is then generally assumed that most of the affected genes will lack expression in the mutant compared with the wildtype with the intact genes. There can, however, be compensatory upregulation in other genes that can hide the full effects of such deletion (18). Since in addition, we also want to investigate the transcriptional effect of the CKb mutations on the genes for the specific proteins found in the previous pulldown and directly interact with CKa as our previous work indicated that these proteins should be helped form the proper interactions by CK2. Compensatory upregulation of the corresponding genes should be expected to compensate for inefficient CK2 function in b1 and b2 mutants if CK2 facilitates IDP forming functional interactions; thus, this is specially investigated.

In this study, transcriptome sequencing showed that the deletion of protein kinase MoCK2 in *M. oryzae* seriously affected cellular metabolism to reduce the acetyl-CoA production. Compared with the background strains Ku80, the expression of genes involved in carbohydrate metabolism, fatty acid metabolism and amino acid metabolism in b1, b2 mutants was down-regulated, then affecting the acetyl-CoA production, which seriously negatively affected the process of energy and substrate metabolism, causing the observed phenotypic defects in the deletion mutants. Several genes encoding proteins found to interact with CKa in the previous pulldown were upregulated. This compensatory upregulation indicates that the CKb1 and 2 are both important for many aspects of the fungal growth and pathogenesis. However, CKb1 is likely more important for pathogenesis than CKb2 that seem more important for fast transcriptional responses.

## Results

### 1. Changes in gene expression due to CKb1 and CKb2 deletions

In this study, we considered the genes with P value less than 0.05 and FoldChange absolute value greater than 2 as significantly different expression genes (DEGs). These are relatively strong criteria because we are mainly interested in “downstream effector genes” with a direct effect on phenotypes since the changes in expression of these genes could explain why we observe the deletion phenotypes in culture. These criteria identified 1297 differentially upregulated genes and 1025 down-regulated genes compared with the background strains Ku80 for b1. For b2 the values were slightly lower with 1189 differentially upregulated genes and 862 down-regulated genes (**Fig.1** and **Table S2**).

**Fig.1.**
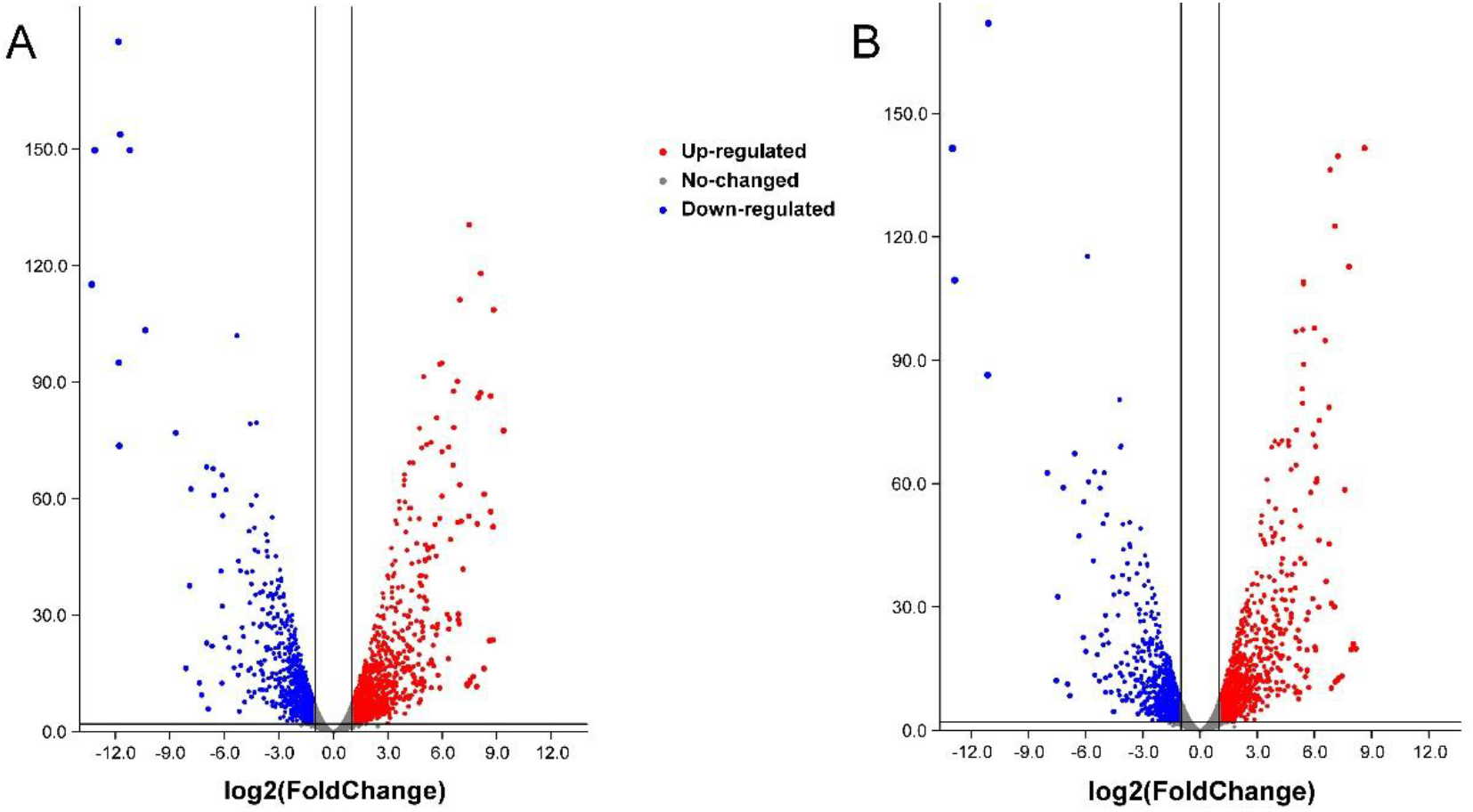
the different expression genes (DEGs) in b1 mutant (A), b2 mutant (B) compared with back ground strains Ku80. The blue points shown the down-expression genes in both mutants compared with back ground strains Ku80. The red points shown the up-expression genes in both mutants compared with back ground strains Ku80. We considered the genes with P value less than 0.05 and FoldChange absolute value greater than 2 as significantly different genes.

In order to understand the function of differential expression genes (DEGs) between b1, b2 and background strains Ku80, KEGG (Kyoto Encyclopedia of Genes and Genomes) (19)and GO (Gene Ontology) were used to analyze the DEGs through the TBtools(20). We focused mainly on the genes down-regulated in the mutant since these are the ones to be most positively affected by an intact CK2 function. KEGG analysis of differential expression genes (DEGs) between b1, b2 and background strains Ku80 showed that deleting the protein kinase MoCK2 components b1 and b2 affected cell metabolism. Differentially upregulated or down-regulated, the affected genes were significantly enriched for genes involved in metabolism, especially for cell carbohydrate metabolism, lipid metabolism, protein (amino acid) metabolism and secondary metabolism (**Fig.2** and **Table S3**). The number of genes with GO annotation of the upregulated or downregulated DEGs were in the b1 mutant 482 and 552, respectively. Similarly, the b2 showed 506, and 395 GO annotated DEGs, respectively. The significantly enriched GO categories also imply involvement in carbohydrate metabolism, lipid metabolism, and protein (amino acid) metabolism, confirming what was found for the KEGG-classified proteins (**Fig.3** and **Table S3, sheet5-12**). From the results of the KEGG and the GO analysis, we could note that protein kinase MoCK2 component deletions affected critical metabolic processes in cells and played an essential role in the growth and development of *M. oryzae* that can result in the obvious phenotypic defects previously observed.

**Fig.2.**
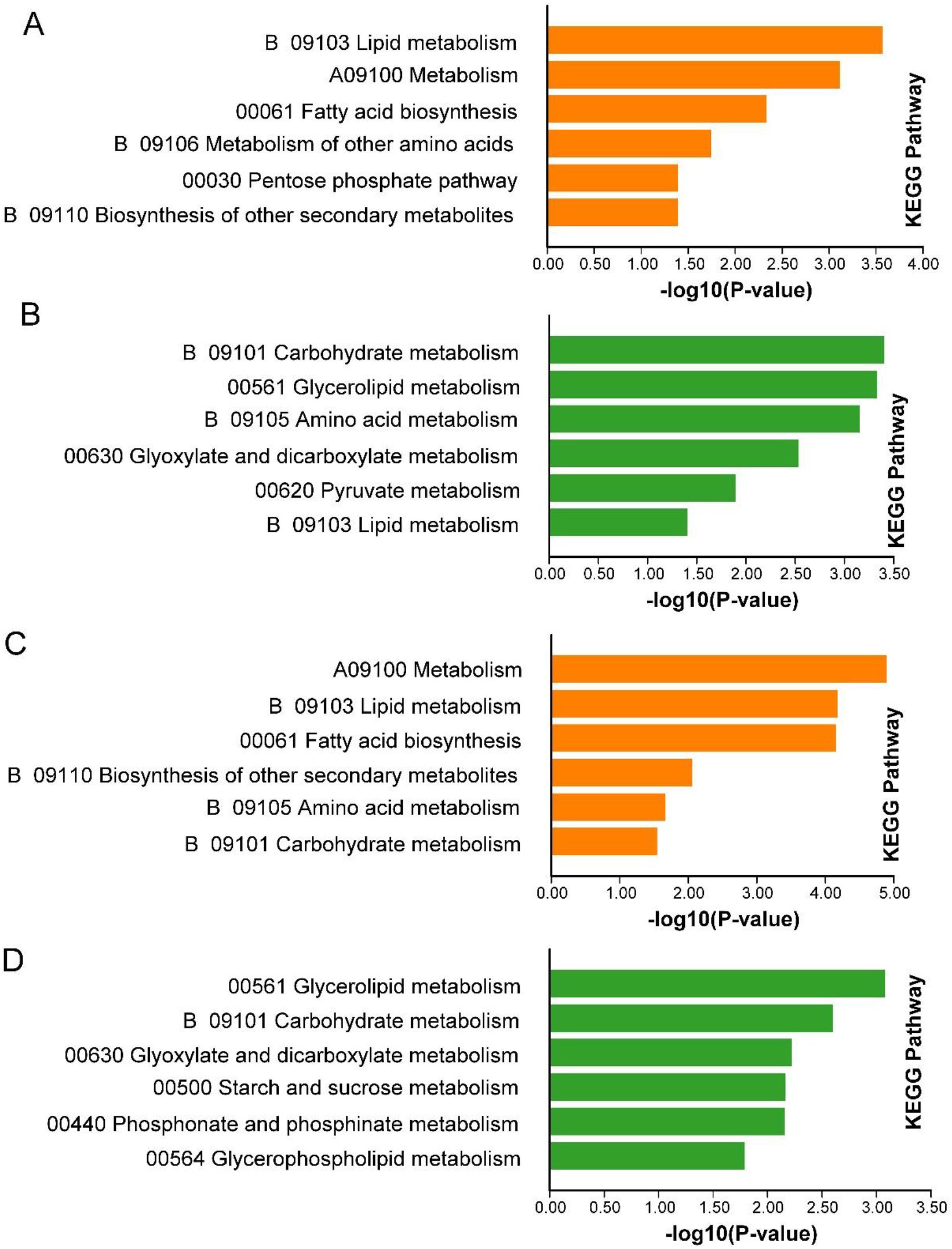
Some results of KEGG enrichment of DEGs in mutants b1, b2 compared with background strains Ku80. A indicates the KEGG enrichment of up-regulated genes in b1 mutant compared with Ku80; B indicates the KEGG enrichment of down-regulated genes in b1 mutant compared with Ku80; C indicates the KEGG enrichment of up-regulated genes in b2 mutant compared with Ku80; D indicates the KEGG enrichment of down-regulated genes in b2 mutant compared with Ku80. The P value of the KEGG enrichment terms showed here are less than 0.05.

**Fig.3.**
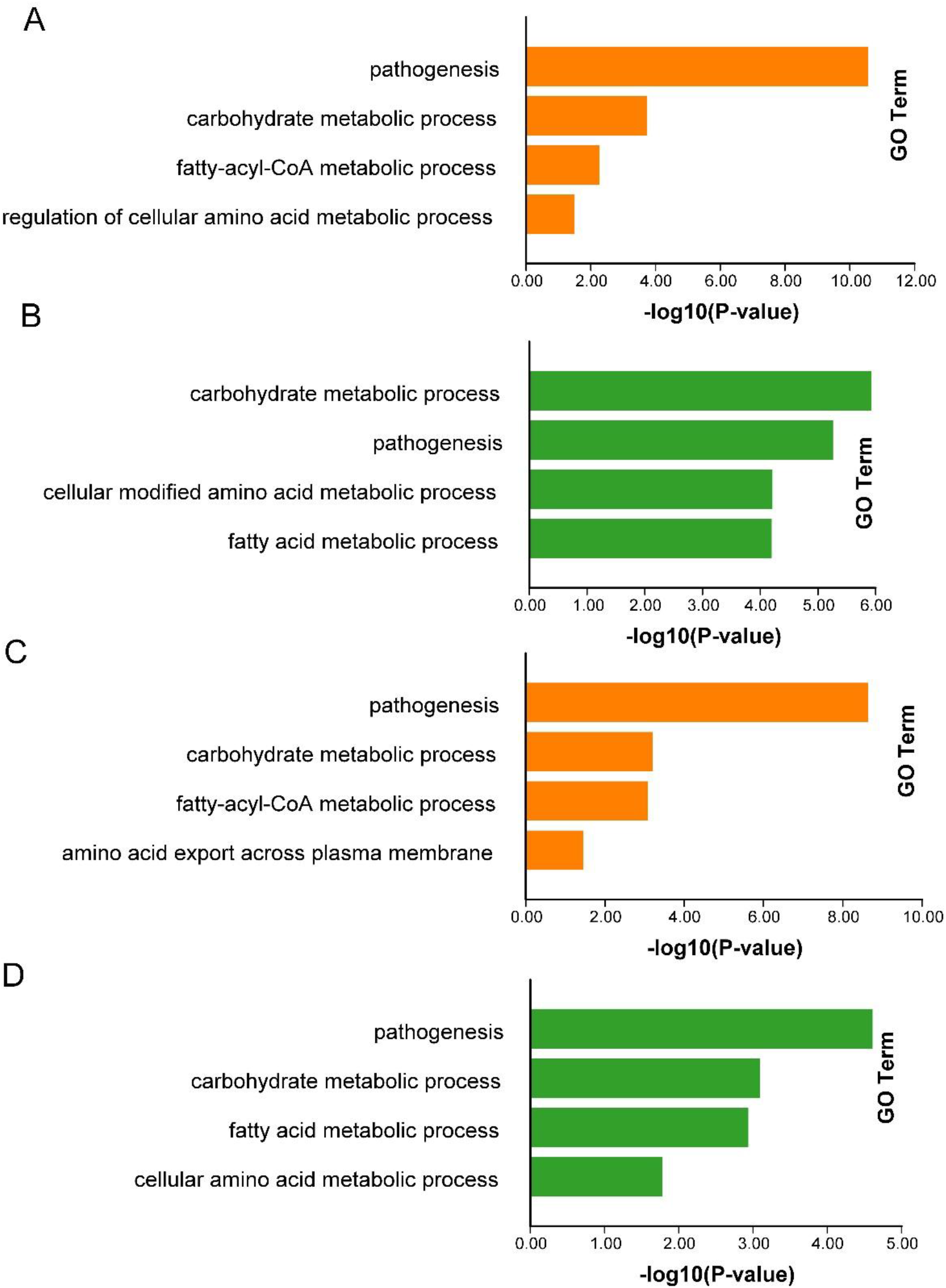
Some results of GO enrichment of DEGs in mutants b1, b2 compared with background strains Ku80. A indicates the GO enrichment of up-regulated genes in b1 mutant compared with Ku80; B indicates the GO enrichment of down-regulated genes in b1 mutant compared with Ku80; C indicates the GO enrichment of up-regulated genes in b2 mutant compared with Ku80; D indicates the GO enrichment of down-regulated genes in b2 mutant compared with Ku80. The P value of the GO enrichment terms showed here are less than 0.05.

### 2. Acetyl-CoA synthesis is affected in the deletion mutants

Acetyl-CoA is a link between various types of metabolisms(21). Therefore, we focused on which of these could be affected by the protein kinase MoCK2.

The primary sources of acetyl-CoA in a cell are the carbohydrate catabolic process, including glycolysis and the dehydrogenation of pyruvate, the fatty acid β-oxidation and amino acid oxidative decomposition. The central role of acetyl-CoA in catabolism is to convert chemical energy through the cyclic oxidation of tricarboxylic acid, or fatty acids, to more generally useful energy intermediates (ATP, NADH, NADPH, Electrochemical membrane potentials) and or participate in anabolic pathways by supplying biosynthesis “building blocks” (21) (**Fig. S3**). Thus, we looked at the change in expression of genes involved in acetyl-CoA metabolism to detect whether MoCK2 could affect acetyl-CoA metabolism.

Firstly, we detected genes annotated as belonging to carbohydrate metabolism in both mutants. The results showed that deleting MoCK2 components likely seriously affected the carbohydrate metabolism in cells. In the b1 mutant, 226 genes with upregulated DEGs have annotation for the carbohydrate metabolism pathway (GO:0005975), and 274 genes with down-regulated DEGs were significantly enriched for genes annotated as GO:0005975(P<=0.05, and, respectively). The number of genes enriched for GO:0005975 accounted for 48% of the DEGs recorded for the b1 mutant. It was similar for the b2 mutant in that there were 234 genes with upregulated DEGs and 188 genes with down-regulated expression significantly enriched for genes annotated as GO:0005975(P<=0.05, and, respectively). The number of genes enriched in GO:0005975 accounted for 46% of DEGs in the b2 mutant (**Fig.4** and **Table S4, sheet1-4**). The genes involved in carbohydrate catabolism were mainly enriched in two GO terms: cellular carbon catabolic process (GO:0044275) and carbon catabolic process (GO:0016052) (**Fig.4A-D** and **Table S4, sheet5,7**). The number of upregulated DEGs from both mutants significantly enriched in GO:0044275 and GO:0016052 was 74 (P<=0.05, and, respectively), and the number of down-regulated DEGs significantly enriched was 66 (P<=0.05, and, respectively) (**Fig.4E, F** and **Table S4, sheet6,8**). Some enzymes in carbohydrate metabolism, such as endoglucanase and glycoside hydrolase, have many coding genes in cells. Their expression changes also show different trends, some have upregulated DEGs, and some are down-regulated, indicating the complexity of CK2 affecting carbohydrate metabolism.

**Fig.4.**
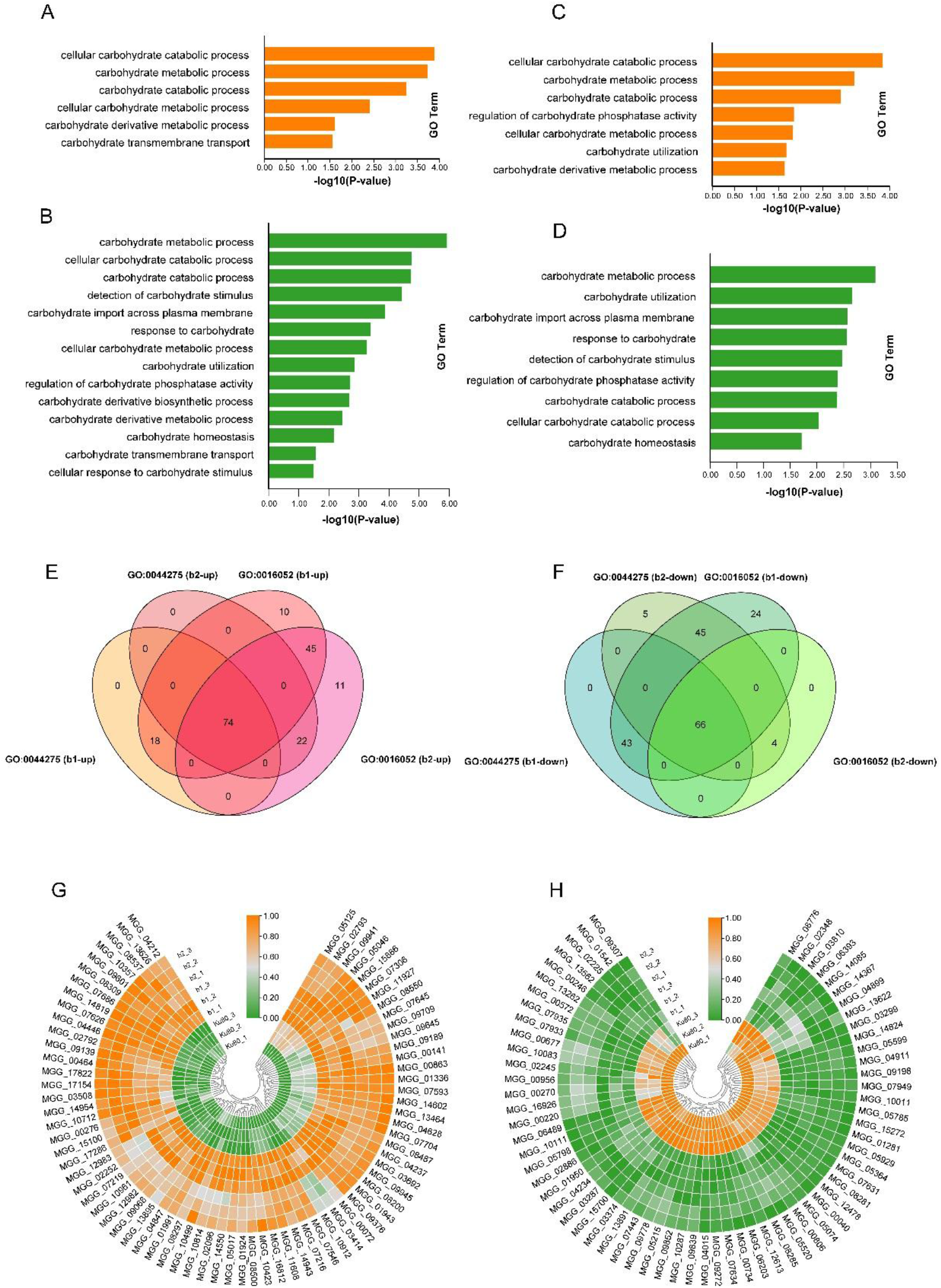
Go enrichment and expression heat map of some genes involving in the carbohydrate metabolic process in b1, b2 mutant. A showed that GO enrichment of up-regulated genes involving in the carbohydrate metabolic process in b1 mutant; B showed that GO enrichment of down-regulated genes involving in the carbohydrate metabolic process in b1 mutant; C showed that GO enrichment of up-regulated genes involving in the carbohydrate metabolic process in b2 mutant; D showed that GO enrichment of down-regulated genes involving in the carbohydrate metabolic process in b2 mutant; E showed the number of up-regulated genes from both b1 and b2 mutants enriched in GO:0044275 and GO:0016052 was 74; F showed the number of down-regulated genes from both b1 and b2 mutants enriched in GO:0044275 and GO:0016052 enriched was 66; G showed the heat map indicating the 74 up-regulated genes enriching in GO:0044275 and GO:0016052 in b1 and b2 mutants; H showed the heat map indicating the 66 down-regulated genes enriching in GO:0044275 and GO:0016052 in b1 and b2 mutants. Genes showing similar patterns of gene expression are clustered.

In addition, the upregulated gene, MGG_10423, encoding cellulase and the down-regulated gene, MGG_03287, encoding amylase, indicated that MoCK2 had diverse regulatory mechanisms for the catabolism of macromolecular sugars in *M. oryzae.* Moreover, some sugar transporters, such as high-affinity glucose transporter, hexose transporter and sugar transporter STL1, had an undeniable down-regulation trend in both mutants, indicating that the deletion of MoCK2 affected the absorption and transport of sugars, resulting in significant changes in the expression of related genes in glucose metabolism, especially catabolism (**Fig.4G, H** and **Table S4, sheet6,8**).

Glucose is converted into pyruvate through glycolysis and mitochondrial import. Pyruvate is in the mitochondria converted to acetyl-CoA through a series of reactions starting with pyruvate dehydrogenase(22). Therefore, we tested whether the expression of the mitochondrial pyruvate dehydrogenase coding gene and the following reactions in the b1 and b2 mutants were affected. The results showed that in b1 and b2 mutants, 11 genes with downregulated DEGs were significantly enriched for mitochondrial pyruvate dehydrogenase metabolism genes (**Fig.5A-C** and **Table S5).** At the same time, we also found that the expression of pyruvate transmembrane transporter genes also showed a downward trend in b1 mutant, indicating that the deletion of MoCK2 could also affect pyruvate transmembrane transport between cytoplasm and mitochondria or peroxisomes (**Fig.5D** and **Table S5**). These results above suggested that MoCK2 deletion affected the conversion of pyruvate to acetyl-CoA, affecting the downstream metabolism.

**Fig.5.**
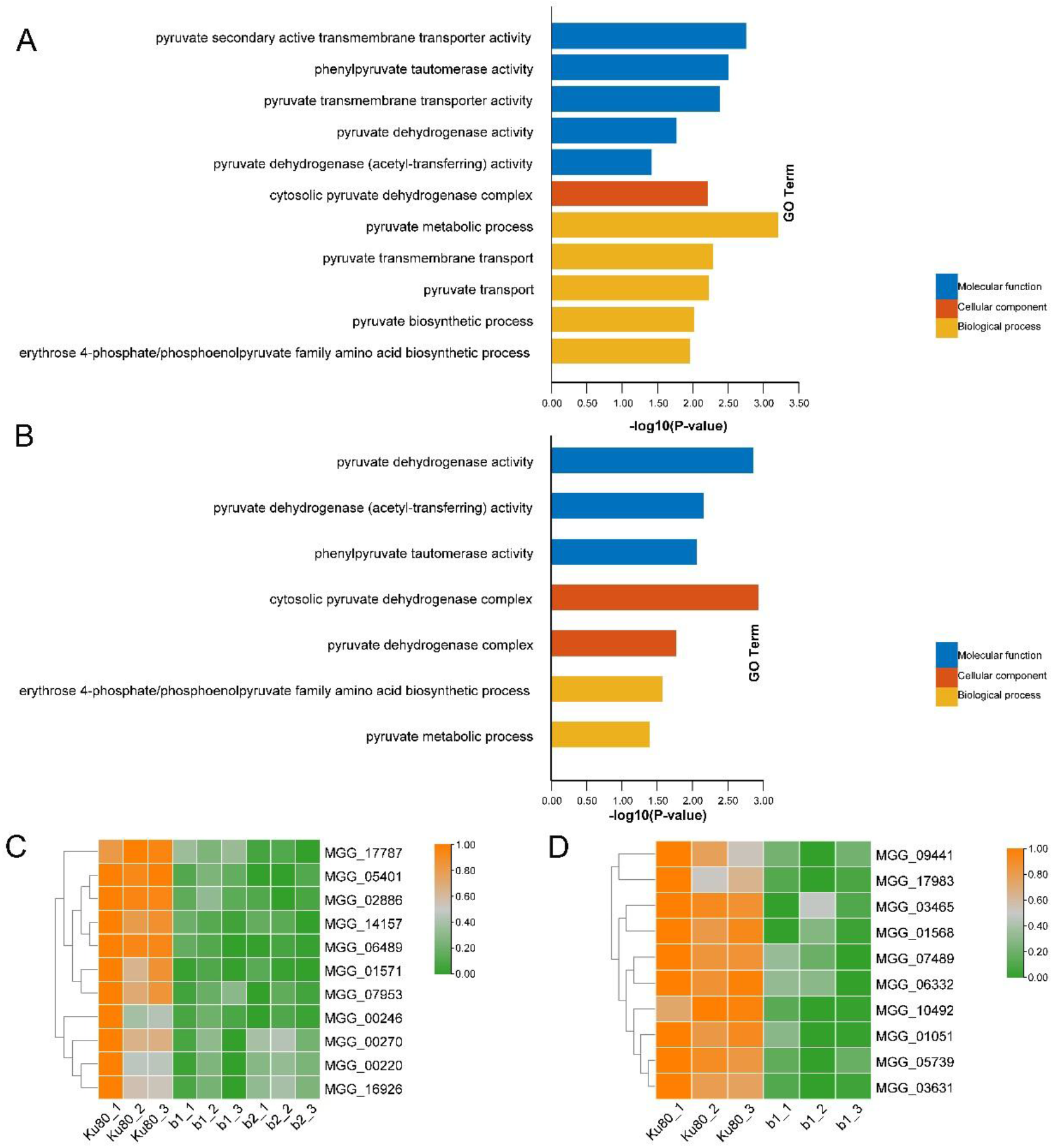
Analysis the genes expression involving pyruvate dehydrogenase in b1, b2 mutant. A showed GO enrichment of pyruvate metabolism in b1 mutant; B showed GO enrichment of pyruvate metabolism in b2 mutant; C indicated the heatmap showing the gene expression of genes encoding pyruvate dehydrogenase in b1, b2 mutants. D indicated the heat map showing the expression of the gene encoding pyruvate transmembrane transporter in b1 mutant. Genes showing similar patterns of gene expression are clustered.

The results above indicate that the deletion of protein kinase MoCK2 reduced the catabolism of carbohydrates and the activity of pyruvate dehydrogenase in cells, and then reduced the content of acetyl-CoA, thus slowing down the metabolism of cells, which was an essential element of the observed phenotypic defects of both mutants(15).

### 3. Acetyl-CoA synthesis from fatty acid metabolism is affected in the deletion mutants

Fatty acid catabolism, especially β-oxidation, is a significant source of acetyl-CoA, especially in conidia (23) (**Fig.6A**), and we detected expression changes of related genes (**Fig.6B, C** and **Table S6, sheet1-4**). The results showed that 42 down-regulated genes were enriched in GO term of fatty acid catabolic process (GO:0009062) in the b2 mutant, 38 downregulated in the b1 mutant. (**Fig.6D, E** and **Table S6, sheet5**). Further analysis showed that two critical enzymes, carnitine acetyltransferase, encoded by MGG_06918, and acyl-CoA dehydrogenase, encoded by MGG_03418, were down-regulated in both deletion mutants. Carnitine acetyltransferase(24), which transports acetyl-CoA between subcellular compartments, controls the beginning of fatty acid oxidation. Acyl-CoA dehydrogenase converts acyl-CoA to enoyl-CoA and reduces FAD to FADH2 (Reduced flavin adenine dinucleotide)(25, 26). FADH2 enters the mitochondrial electron transport chain for oxidation and participates in the generation of the pH gradient over the mitochondrial inner membrane, resulting in ATP exported to the rest of the cell and exchanged for ADP. The reduced expression of carnitine acetyltransferase and acyl-CoA dehydrogenase likely affects the oxidative degradation of fatty acids, reducing the cellular content of acetyl-CoA and affecting the energy conversion metabolism resulting in that cells do not have enough available energy in suitable forms for driving cellular anabolic processes. In addition, fatty acid transmembrane transporters were enriched (GO:1902001). The number of these genes affected in both mutants was 22 (**Fig.6F, G** and **Table S6, sheet6**). The decrease in the expression of transporter coding genes indicates that the deletion of protein kinase MoCK2 reduced the transmembrane transport of fatty acids in cells, which can directly affect the rate of fatty acid oxidation.

**Fig.6.**
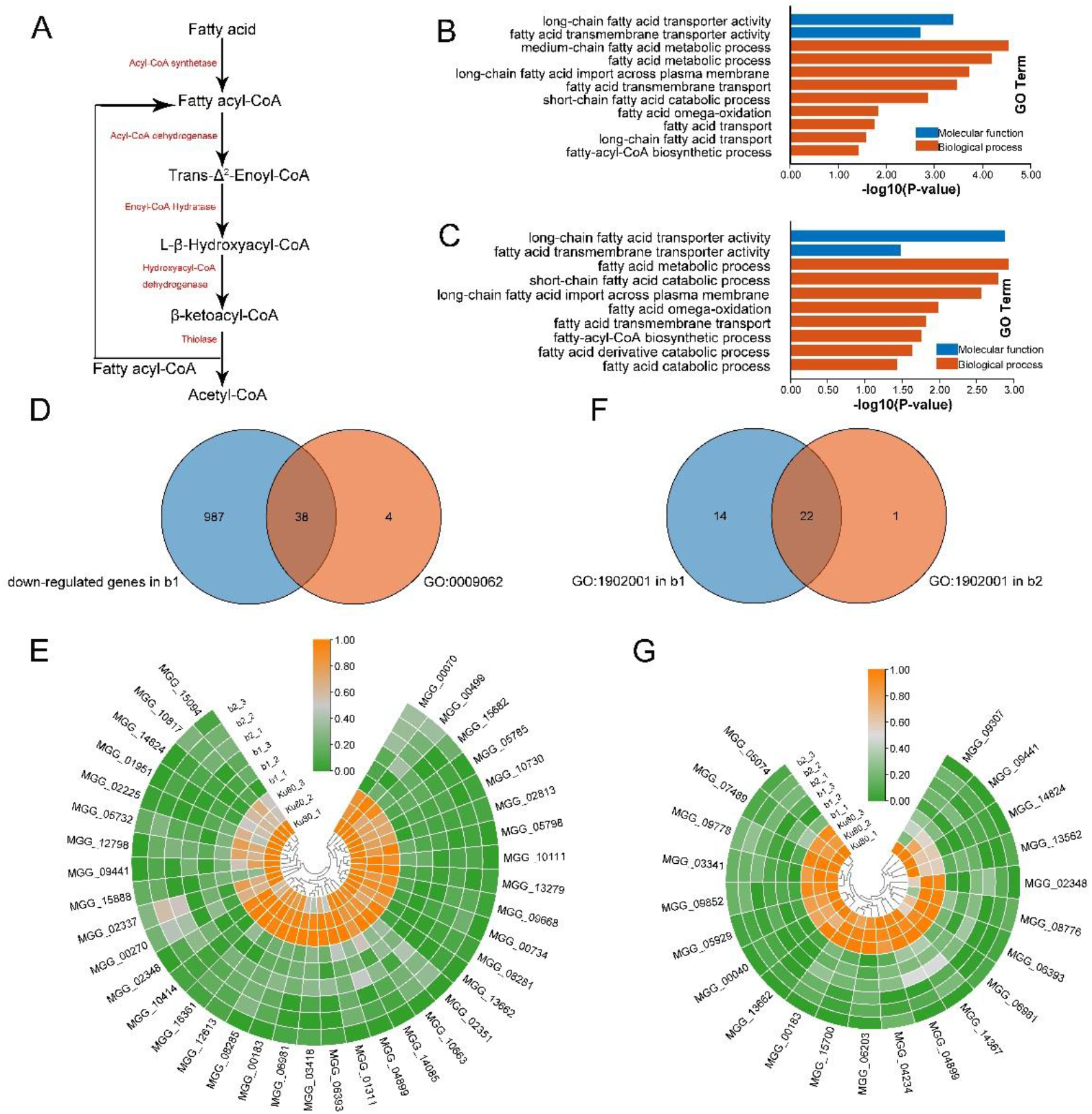
Analysis the genes expression involving fatty acid metabolism in b1, b2 mutant. A showed the fatty acid beta-oxidation pathway. B and C showed DEGs GO enrichment in fatty acid metabolism in b1, b2 mutant, respectively. D: Venn plot indicated 38 genes enriched in fatty acid catabolic process (GO:0009062) in b1, b2 mutants. E showed the expression of 38 genes enriched in fatty acid catabolic process (GO:0009062) in b1, b2 mutants compared with background strains Ku80. MGG_06918 encodes carnitine acetyltransferase; MGG_03418 encodes acyl-CoA dehydrogenase. F: Venn plot indicated 22 genes enriched in fatty acid transmembrane transport (GO:1902001) in b1, b2 mutants. G showed the expression of 22 genes enriched in fatty acid transmembrane transport (GO:1902001) in b1, b2 mutants compared with background strains Ku80. Genes showing similar patterns of gene expression are clustered.

In conclusion, the deletion of MoCK2 likely affected the production of acetyl-CoA by reducing fatty acid metabolism and intracellular transport. It then affected the subsequent formation rate of cellular energy “currencies” as ATP needed for growth, maintenance and synthesis of extracellular compounds, resulting in the previously observed severe phenotypic defects (15).

### 4. The acetyl-CoA synthesis from amino acid metabolism is affected in the deletion mutants

For example, amino acid catabolism, alanine, glycine, serine, threonine and phenylalanine are other significant sources of acetyl-CoA. Among the down-regulated genes in both mutants, we found GO enrichment terms of amino acid catabolism (**Fig.7A, B** and **Table S7, sheet1-4**). GO:0009063 represent the cellular amino acid catabolic process, in which there were 88 genes downregulated in b1 mutant and 58 genes in b2 mutant. 50 of these genes were down-regulated in both mutants (**Fig.7C, D** and **Table S7, sheet5**). The deletion of any MoCKb seriously inhibited the catabolism of amino acids. In addition, we found the aspartate family amino acid catabolic process (GO:0009068), serine family amino acid catabolic process (GO:0009071), and glutamine family amino acid catabolic process (GO:0009065) are downregulated in the b1 mutant (**Fig.7E-G** and **Table S7, sheet6-8**). The catabolites of aspartate and glutamine to form α-ketoglutarate and oxaloacetate can directly participate in the tricarboxylic acid cycle. Serine catabolism is another source of acetyl-CoA.

**Fig.7.**
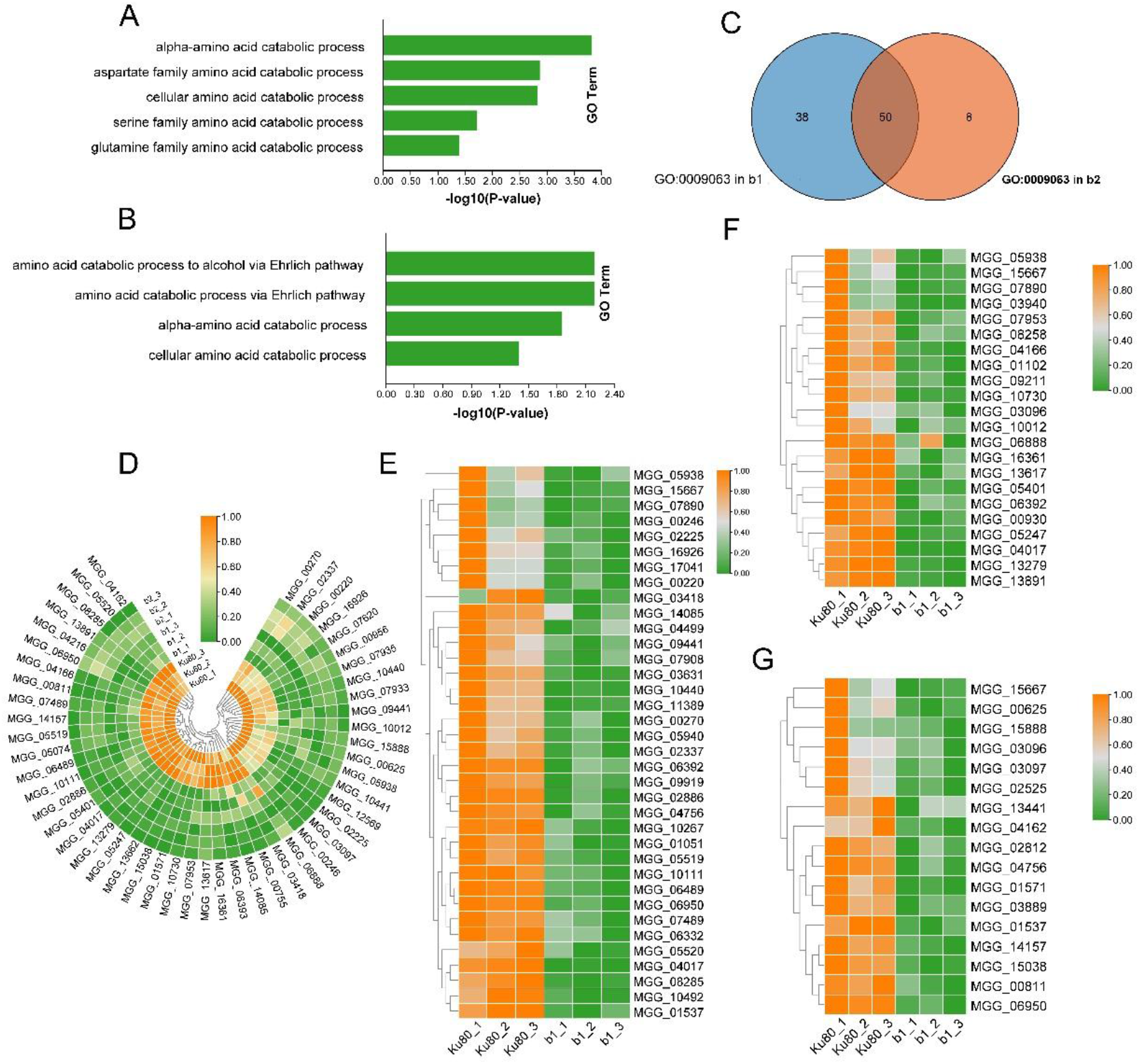
the expression of genes involving in the amino acid catabolic pathway. A showed the GO enrichment of amino acid catabolic process in b1 mutant; B showed the GO enrichment of amino acid catabolic process in b2 mutant; C showed that the genes enriched in GO:0009603 form both b1, b2 mutant; D: the heatmap showed the expression of 50 genes enriching in GO:0009603 form both b1, b2 mutant; E: the heatmap showed the expression of genes enriching in aspartate family amino acid catabolic process (GO:0009068) in b1 mutant; F: the heatmap showed the expression of genes enriching in glutamine family amino acid catabolic process (GO:0009065) in b1 mutant; G: the heatmap showed the expression of genes enriching in serine family amino acid catabolic process (GO:0009071) in b1 mutant. Genes showing similar patterns of gene expression are clustered.

### 5. Expression of transporter encoding genes is affected in the deletion mutants

In the preface, we have simply introduced that the deletion of MoCK2 did affect the transmembrane transport of carbohydrate transporters, pyruvate and fatty acids. So, we suspected that the blocked transport of various substances was an important reason for the phenotypic defects of b1, b2 mutants.

We found that the expression of the gene encoding carbohydrate transporter was severely inhibited in the mutant. The expression of 11 genes was down-regulated in both b1, and b2 mutants. These genes were MGG_00040 and MGG_06203 encoding high-affinity glucose transporter, MGG_05929, MGG_04234 and MGG_09307 encoding hexose transporter, MGG_09852 encoding sugar transporter STL1, MGG_03341 encoding plastidic glucose transporter 4, MGG_02394 and MGG_05116 encoding malic acid transporter, MGG_09441 encoding tricarboxylate transporter, MGG_04371 encoding general alpha-glucoside permease (**Fig.8A** and **Table S8, sheet2,4**). In addition, b1 knockout affected the absorption and transport of lactose and maltose because of the down-regulation of MGG_05889, MGG_09228, MGG_15388 encoding lactose permease and MGG_08266 encoding maltose permease MAL61 (**Fig. 8C** and **Table S8, sheet2**). These results showed that the absence of protein kinase MoCK2 decreased the ability of mycelium to absorb and utilize sugar from the medium.

**Fig.8.**
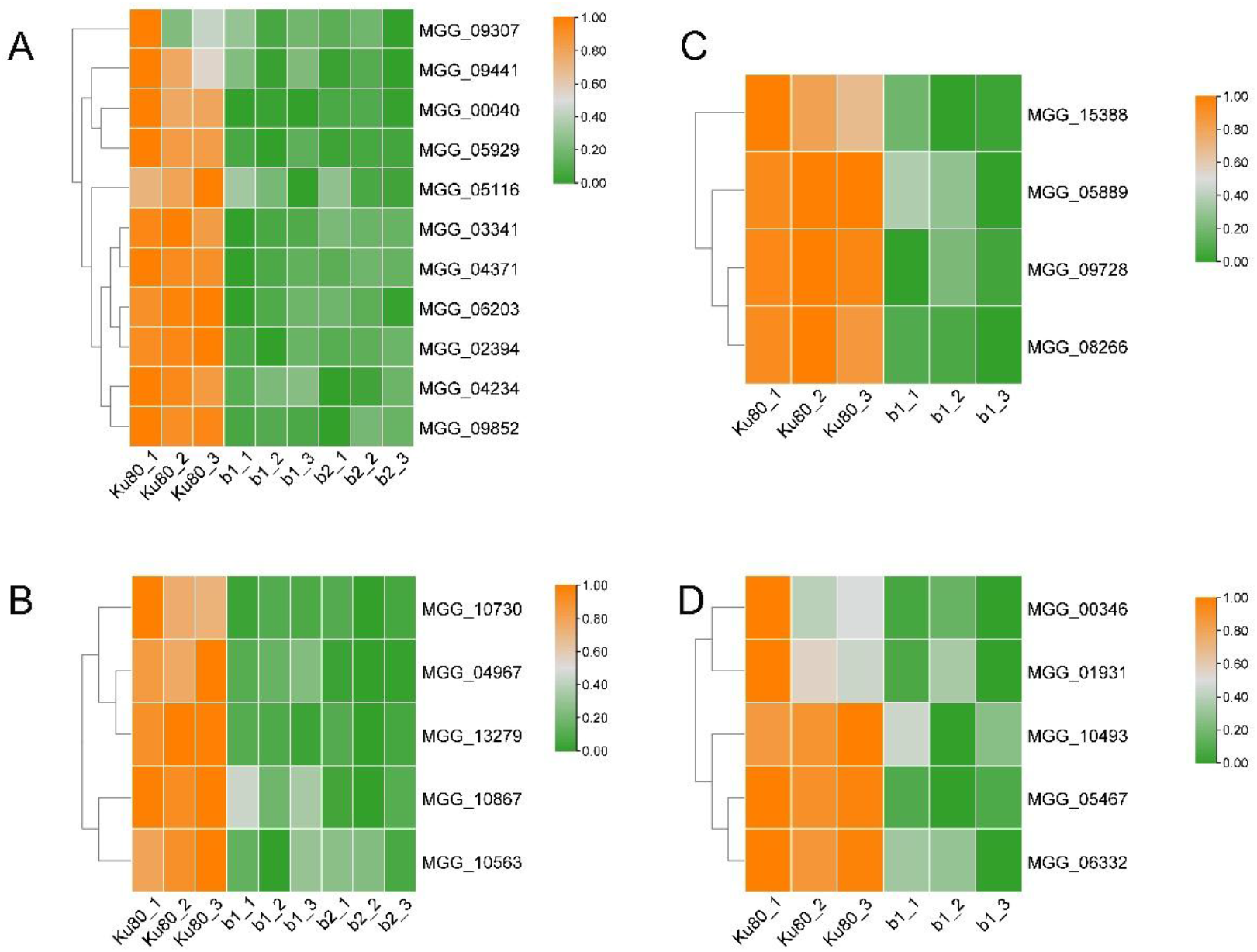
the expression of genes encoding transporter in b1, b2 mutant compared with background strains Ku80. A indicated the expression of genes encoding carbohydrate transporters in both b1, b2 mutant compared with Ku80; B indicated the expression of genes encoding metal ion transporters in both b1, b2 mutant compared with Ku80; C indicated the expression of genes encoding lactose permease and maltose permease in b1 mutant compared with Ku80; D indicated the expression of genes encoding metal ion transporters in b1 mutant compared with Ku80. Genes showing similar patterns of gene expression are clustered.

Besides free diffusion, the absorption of sugars is accompanied by an active transport mode of ion diffusion. Meanwhile, we also found that the expression of some ion transporter coding genes was down-regulated in mutants, such as MGG_05467 encoding sodium/nucleoside cotransporter 1, MGG_06332 encoding peroxisomal adenine nucleotide transporter 1, MGG_10867, MGG_04967 and MGG_10493encoding zinc-regulated transporter, MGG_00346 encoding pi-transporter, MGG_01931 encoding siderophore iron transporter mirB (**Fig.8B, D** and **Table S8, sheet2,4**).

Compared with the downregulation of glucose transporters, the expression trend of genes encoding amino acid transporters in both mutants was not particularly obvious. Regulated amino acid transporters were found among the mutant’s up- and down-regulated genes. MGG_08426, MGG_14203 and MGG_05833 encoding amino acid transporter were down-regulated in both mutants (**Table S8, sheet2,4**). MGG_00289 encoding amino-acid permease inda1 was down-regulated. However, MGG_11327 and MGG_06036, which also encode amino-acid permeases, were upregulated in the b1 mutant (**Table S8, sheet1,2**). Similarly, proline permease encoding genes MGG_04216 and MGG_14937 were downregulated in the b1 mutant, but MGG_08548 was upregulated in the b2 mutant (**Table S8**). These results show that amino acid transporters had complex regulation in response to CK2 deletion.

### 6. Genes known to directly interact with CKa are upregulated in the CKb mutants

Of the significantly upregulated genes in the b1 and b2 mutants, 59 of the encoded proteins were found in the previous CKa pulldown (**Table S9** and **S10**). Since CKa is expected to phosphorylate these proteins and have a chaperone effect(16), a less efficient CK2 holoenzyme should chaperone these proteins and lead to compensatory upregulation of their corresponding genes. Our results show that this is indeed the case (**Fig.9**), where it is shown that genes for proteins directly interreacting with CKa are upregulated more, the more it is found in the CKa pulldown. The relationship between these two measures is the same for the 60 significantly upregulated genes b1 and b2 in that the two slopes for the fitted lines of the correlations are the same (**Fig.10**). Interestingly, some significantly regulated genes for b1 (8) and b2 (9) are only upregulated in one of the mutants indicating a differential involvement for CKb1 and CKb2 in regulating the phosphorylation by CKa in the CK2 holoenzyme (**Fig.10, Fig.S4**).

**Fig.9.**
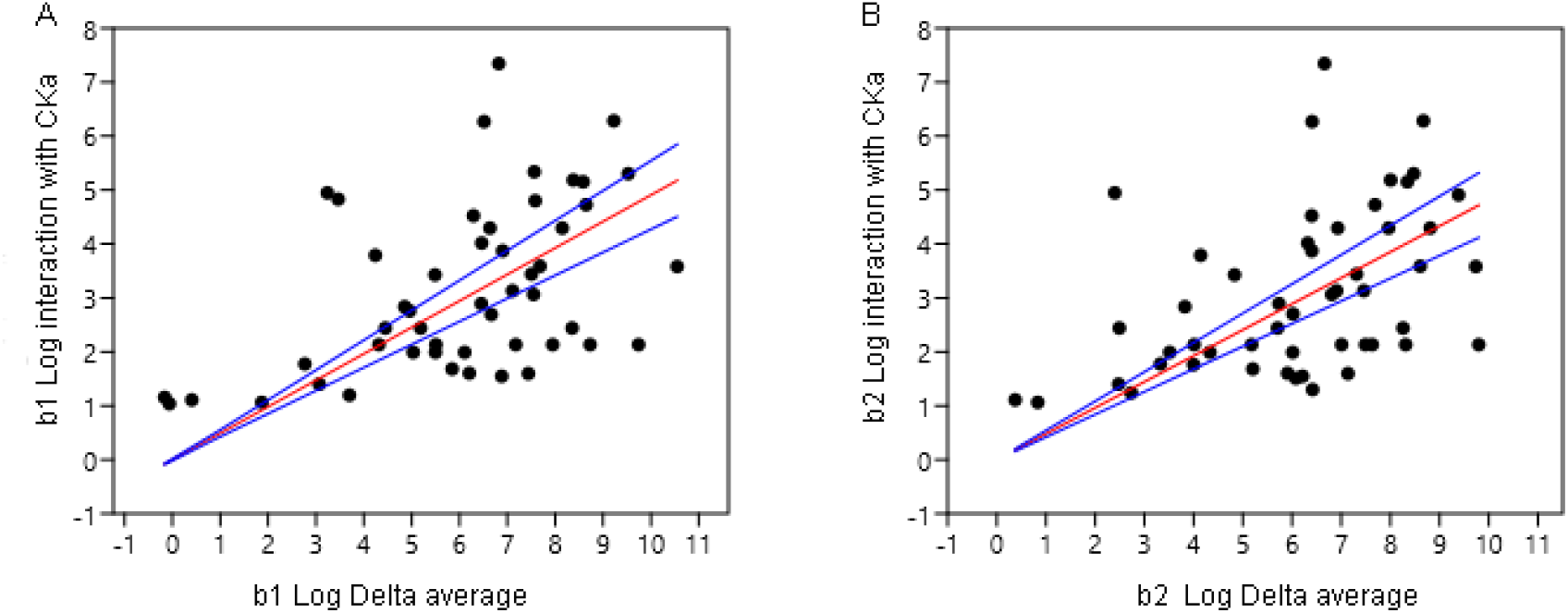
Upregulated genes also found in the CKa pulldown in a previous paper are correlated with the amount of their encoded proteins found in the CKa pulldown. A shows that over a threshold the upregulation of genes in the mutant b1 for proteins found in the CKa pulldown is in general correlated with the amount of proteins of respective proteins found in the pulldown (blue lines=+-95% conf for red line) P=not correlated=0.00044). B shows a similar plot as in A but for the b2 mutant (blue lines=+-95% conf for red line) P=not correlated=0.000849.

**Fig.10.**
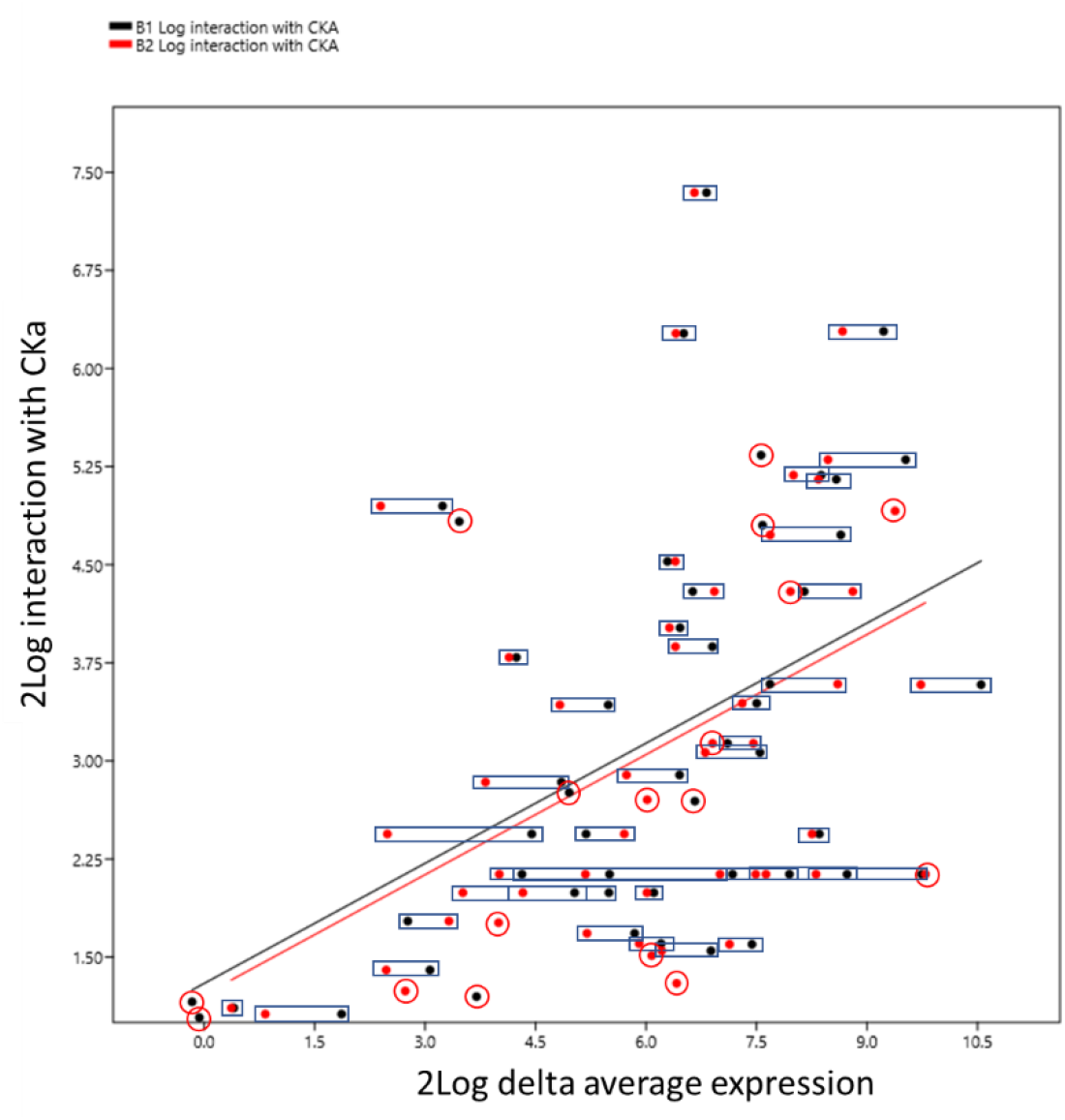
Combined plot of the Fig.9A and B data to highlight that the two relationships shown in Fig.9 are very similar but with some genes upregulated in only one mutant. Black dots are b1 mutant average regulation. Red dots are b2 mutant average regulation. Black rectangles join regulations of the same gene in b1 and b2. Red circles surrounds dots for genes only significantly upregulated in one of the mutants.

Genes for gene products interacting with CKa compensatory upregulated in both the b1 and the b2 mutant were significantly enriched for genes involved in many categories important for plant infection and resistance to plant defences (**Fig.11A** and **Fig.S5**). The b1 is enriched explicitly for gene categories of importance for plant pathogenesis (**Fig.11B** and **Fig.S6**), and b2 is significantly enriched for genes necessary for translation (ribosome biogenesis) but also for making non-ribosomal peptides like siderophores needed for iron uptake (**Fig.11C** and **Fig.S7**).

**Fig.11.**
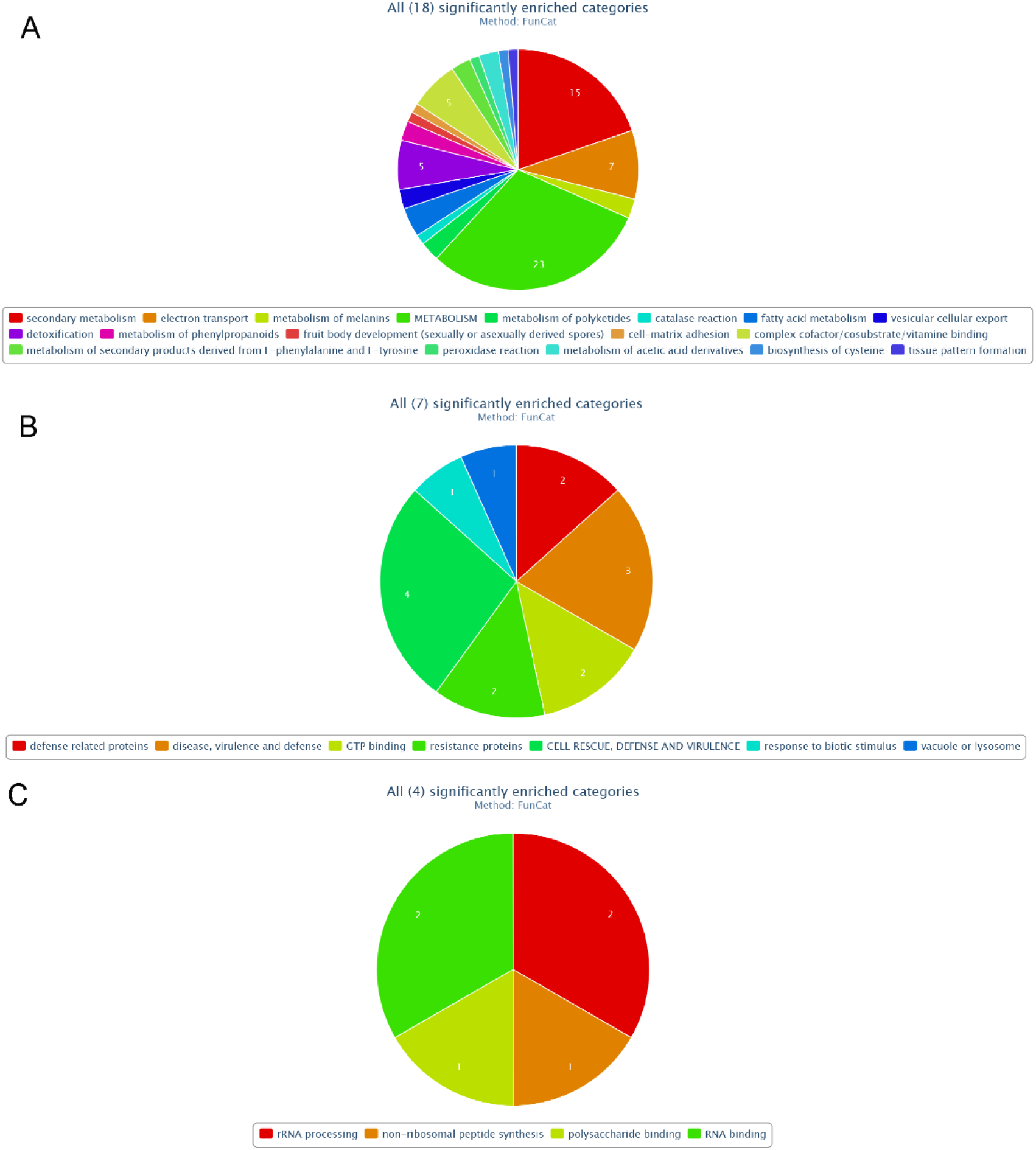
Funcat significantly enriched categories for significantly upregulated genes A. In both b1 and b2 mutants for genes encoding proteins found in a CKa pulldown. B. Only in the b1 mutant. C. only in the b2 mutant.

## Discussion

In previous research, we identified a vital protein kinase MoCK2 which has a critical regulatory function for the growth, sporulation and pathogenicity in *M. oryzae*(15). Moreover, MoCK2 pulldown experiment showed more than 1300 hypothetical substrates(16). Therefore, we used RNA-seq technology to analyze the transcriptome of background strains Ku80 and two mutants b1, b2 to analyze the MoCK2-regulating pathogenic mechanism in *M. oryzae*.

*M. oryzae* must absorb nutrients from the culture medium to maintain life activities. We collected the mycelia of strains Ku80 and b1, b2 cultured in liquid CM medium for transcriptome sequencing. Like growing on a solid medium, the growth rate of b1, and b2 mutants in liquid culture was slower than Ku80. Therefore, we speculate that the deletion of MoCK2 may seriously inhibit the metabolism of *M. oryzae.*

The transcriptome analysis showed that an impaired CK2 holoenzyme caused significant expression changes in many genes in both mutants. KEGG and GO analysis showed that the genes with changed expression were mainly enriched in intracellular metabolisms, such as carbohydrate metabolism, fatty acid metabolism and amino acid metabolism. Further analysis showed that the deletion of MoCK2 seriously hindered the catabolism of substances in cells, directly reducing the concentration of acetyl-CoA in cells.

Acetyl-CoA is an important intermediate in various metabolisms. The impairing of MoCK2 directly appears to slow down pyruvate dehydrogenation and affects the conversion of pyruvate to acetyl-CoA. Catabolism of fatty acids is in yeast, and *M. oryzae* can be carried out in the peroxisomes and acetyl-CoA is initially converted to acetyl-carnitine. Two carnitine acetyl transferases Crat1 and Crat2 have been identified in *M. oryzae,* and deletion of especially Crat1 has large effects on appressorium cell wall formation, utilization of fats and pathogenicity(27). Both Crat1 (MGG_01721) and Crat2 (MGG_06981) were significantly downregulated in the mutation b1 and b2, although around the 2log −1 threshold level. For b1 and b2 Crat1 was found to be −0.81 and −0.56 respectively (P no regulation = 3.33E-05 and 3.14E-03 respectively). For b1 and b2 Crat2 was found to be −1.28 and −1.46 respectively (P no regulation = 1.92E-08 and 4.06E-11 respectively). That downregulation in the CKb mutants can partly explain the previous found phenotypic defects(15).

Thus, fatty acid β-oxidation and amino acid decomposition might be simultaneously reduced by the CK2 deletions, as the expression of related genes was significantly down-regulated. The decreased expression of both Crat1 and Crat2 encoding carnitine acetyltransferase and MGG_03418 encoding acyl CoA dehydrogenase should affect the β-oxidation and reduce the metabolism of acetyl-CoA. Amino acid degradation, especially serine degradation, is also an essential source of acetyl-CoA. Similarly, the expression of genes related to serine degradation in mutants decreased significantly. In addition, the coding genes of a series of transporters involved in metabolism were down-regulated, likely affecting the absorption and transport of sugar, pyruvic acid and fatty acid, further slowing down the corresponding metabolic processes. These results fully demonstrate that MoCK2 participates in the metabolism of acetyl-CoA, as has also been shown for other organisms(28).

CKa interacts with CKb1 and CKb2 to form holoenzymes, and this interaction was confirmed in a pulldown experiment that also showed direct interaction with a large number of other proteins. We concluded in the previous study that CK2 phosphorylation with dephosphorylations likely contributed to the formation of protein-protein and protein-nucleic acid interactions through a chaperone-like mechanism depending on the CK2 phosphorylation(16). The lack of efficiency of the CK2 holoenzyme should consequently lead to a correlation between the absolute upregulation of the proteins and the protein abundance in the previous CKa pulldown, and this was indeed the case (**Fig. 9-10**). The genes compensatory regulated are genes whose gene products should be lacking to be present in the right shape and interaction. Interestingly these genes were overrepresented for genes of importance for pathogenicity (**Fig. 9, Fig. S5-7**) and can further explain the b1 and b2 phenotypes found before(15). Of those genes, many are predicted to encode mitochondrial proteins dependent on chaperone interactions to become imported into the mitochondria, and those were overrepresented among the CKa-interreacting proteins(16). Such destabilization should favour efficient mitochondrial protein import to the mitochondrion through the TOM and TIM complexes (29). In the absence of efficient import, these proteins would need to be compensatorily upregulated, which they are in both b1 and b2 (**Fig. S8**). These proteins were predicted as mitochondrial using DeepMito (30). The predicted genes encoding proteins likely to be mitochondrial (**Table S9**), and present in the previous CKa pulldown, are predicted to locate to the mitochondrial matrix (13), inner membrane (4) and outer membrane (3). There were more of these compensatorily regulated mitochondrial genes in b1 (19 of 20 total, with 5 only in b1) than in b2 (15 of 20 total, with 1 only in b2), further strengthening that the b1 mutant is more severely affected than b2, and in this case for genes encoding mitochondrial located proteins.

Previous studies show that protein kinase MoCK2 accumulates at the mycelial septal pore, in the appressorium, and at the appressorium pore, possibly affecting cellular activities and material exchange between compartments. In addition, MoCK2 also accumulates in the nucleus, where CK2 can affect gene expression and transcription and the formation of ribosomes and other intrinsically disordered protein interactions (15). Such multiple functions were confirmed by the many changes in gene expressions in both mutants. Our previous research obtained many nuclear localization proteins interacting with MoCK2 through a pulldown experiment (16). Therefore, subsequent experiments will further focus on how MoCK2 affects the nuclear-localized proteins in response to the physiological changes and during the rice’s pathogenicity and on the mitochondrial import proteins.

## Material and methods

### Fungal strains and growth conditions

All strains used in this study (background strains Ku80, b1 and b2 mutant) were stored on dry sterile filters paper and cultured in the shaking, complete medium (CM: 0.6% casein hydrolysate, 0.6% yeast extract, 1% sucrose, 1.5% agar) at 25 °C, 200 rpm and harvested by filtering through 3 layers of Miracloth (EMD Biosciences), washed and frozen in liquid nitrogen.

### RNA Extraction and RNA-sequencing

Total RNA was extracted from mycelia using the Plant/Fungi RNA Purification Kit from Sigma-AldrichTrading Co. Ltd (Shanghai, China). We used the Germany IMPLEN P330 ultra-micro spectrophotometer to check the integrity and quantity of RNA. RNA with high integrity was used for library preparations. RNA was prepared from three biological replicates and used for independent library preparations.

High-throughput sequencing was performed on an IlluminaHiseq 2000 machine at IGENEBOOK BIOTECHNOLOGY CO., LTD (Wuhan, China).

### Quality analysis of transcriptome data

The genome of *M. oryzae* was downloaded from the Ensembl Fungi (https://fungi.ensembl.org/). Clean reads were obtained from the raw reads with Trimmomatic, deleting low-quality sequences. The clean RNA-seq reads were aligned to *M. oryzae* genome using hisat2 software (version: 2.0.1-beta)(31). The alignment quality was calculated at the Q20, Q30 and GC content.

Nine sample hyphae for sequencing were collected from the background strains Ku80, b1, b2 mutants. The transcriptome of 9 samples was reconstructed by Stringtie (version: 2.0.4) and Gffcompare software(32, 33). The sequencing platform used was Illunima hiseqtm2000. The data obtained by transcriptome sequencing of nine samples was between 7.1 Gb and 9.5 Gb. After filtering low-quality reads and adaptor sequences, 6.9 Gb~9.4 Gb of high-quality clean reads remained. The Q20 of clean reads ranges from 98.29% to 98.44%, and the Q30 ranges from 94.97% to 95.23%. Besides, the average content of GC is about 56.53%, which was slightly higher than that of the reported genome of *M. oryzae* (**Table S1, sheet1**). The clean RNA-seq reads were aligned to the *M. oryzae* genome using hisat2 software(31). For the nine samples, the results showed that the lowest proportion of mapped reads was more than 83.27% (the highest was 93.64%) in clean reads. 98% of mapped reads were unique (**Table S1, sheet2**).

We counted the effective reads compared to the genome according to the functional elements. We divided the genome into CDS, 5’UTR, 3’UTR, intron and intergenic regions, and counted the proportion of reads belonging to them, respectively. The results showed that more than 77% of the effective reads in 9 samples covered the CDS region, which met the requirements for data analysis (**FigS1** and **Table S1, sheet3**). The transcriptome data of 9 samples were analyzed by Stringtie(32) and Gffcompare(33) software, and compared with the reference genome data, 3023 new genes and 13184 annotated genes were obtained (**Table S1, sheet4** and **sheet5**).

### Analysis of different expressions of genes among 9 samples

After obtaining effective reads, we used the featurecounts software (version: v1.6.0)(34) to count the number of reads on the gene according to the annotation file of the *M. oryzae* genome. Because the sequencing amount of each sample is different, to horizontally compare the expression difference of the same gene among different samples, we needed to standardize the number of reads of the gene. The standardized method was FPKM (fragments per kilobase of exon per million reads mapped)(35). This result is shown in **Table S1, sheet6**. Principal component analysis (PCA)(36), which illustrates the intrinsic biological variation among samples, showed that samples from the same strains grouped (**Fig.S2A**). The heat map, carried out with the heatmap2 function in the gplots package, showed the similarity of gene expression patterns within the same strain, but there were significant differences between the background strain Ku80 and the two mutants (**Fig.S2B**).

In order to compare the difference in gene expression among different samples, we performed differential expression analysis by R-package edgeR(37). As a general rule, we considered genes with P value less than 0.05 and FoldChange absolute value greater than 2 as significantly differentially expressed genes. These different expression genes (DEGs) are shown in **Table S2**.

### Functional enrichment analysis

In this paper, we used the TBtools to do GO annotation, enrichment and KEGG enrichment of differentially expressed genes (DEGs) between b1, b2 mutants and Ku80(19, 20). In addition, we used FungiFun to do FunCat and GO enrichment analysis of genes both upregulated and present in a previous CKa pulldown experiment since for that experiment that webtool had been used for the functional classification of the pulldown proteins. The transcription abundance values of all genes were used to calculate the Euclidean distance between each sample in the heat map analysis.

## Supplementary information

**Fig. S1.**
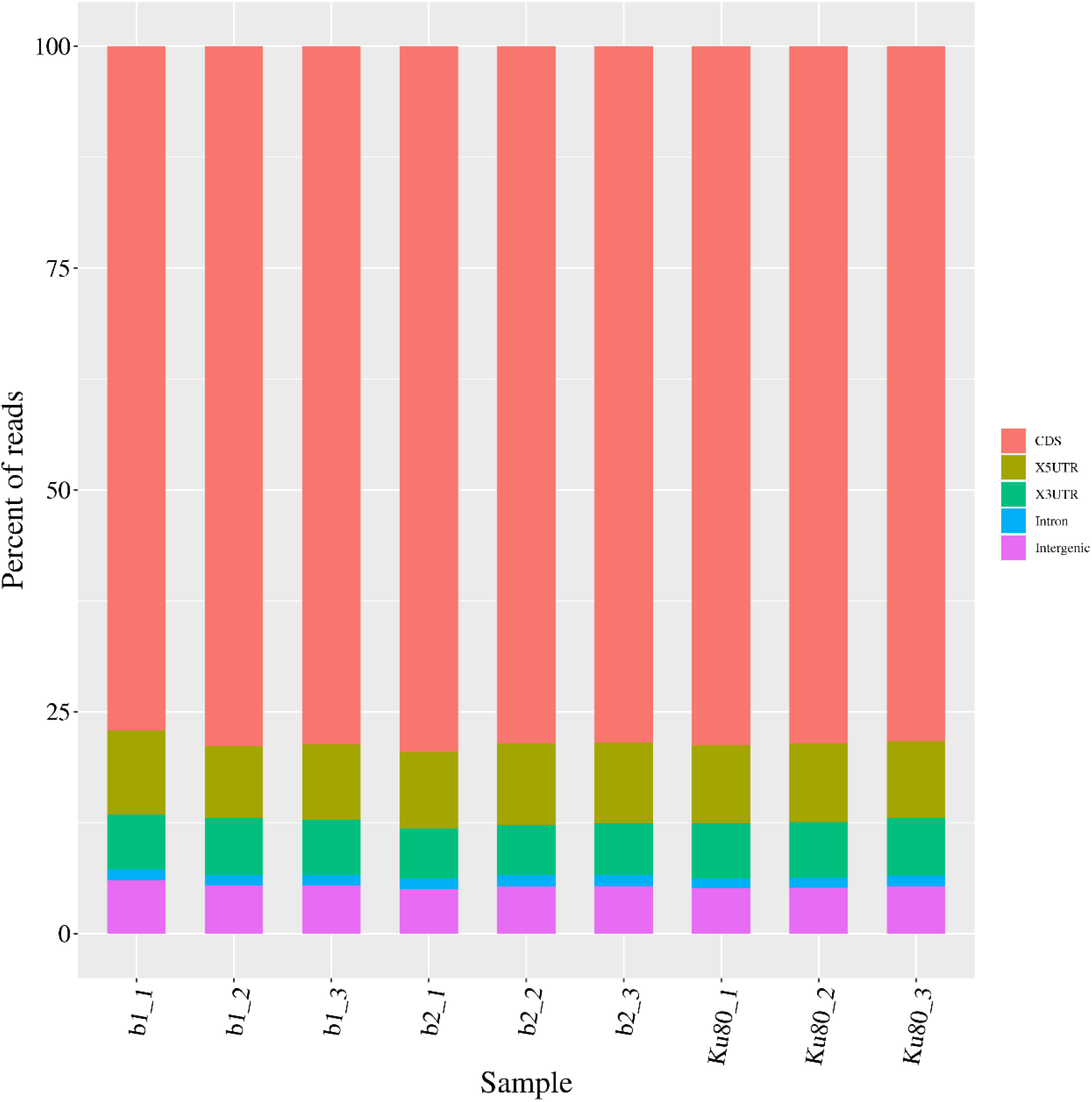
Distribution table of reads on functional gene elements.

**Fig. S2.**
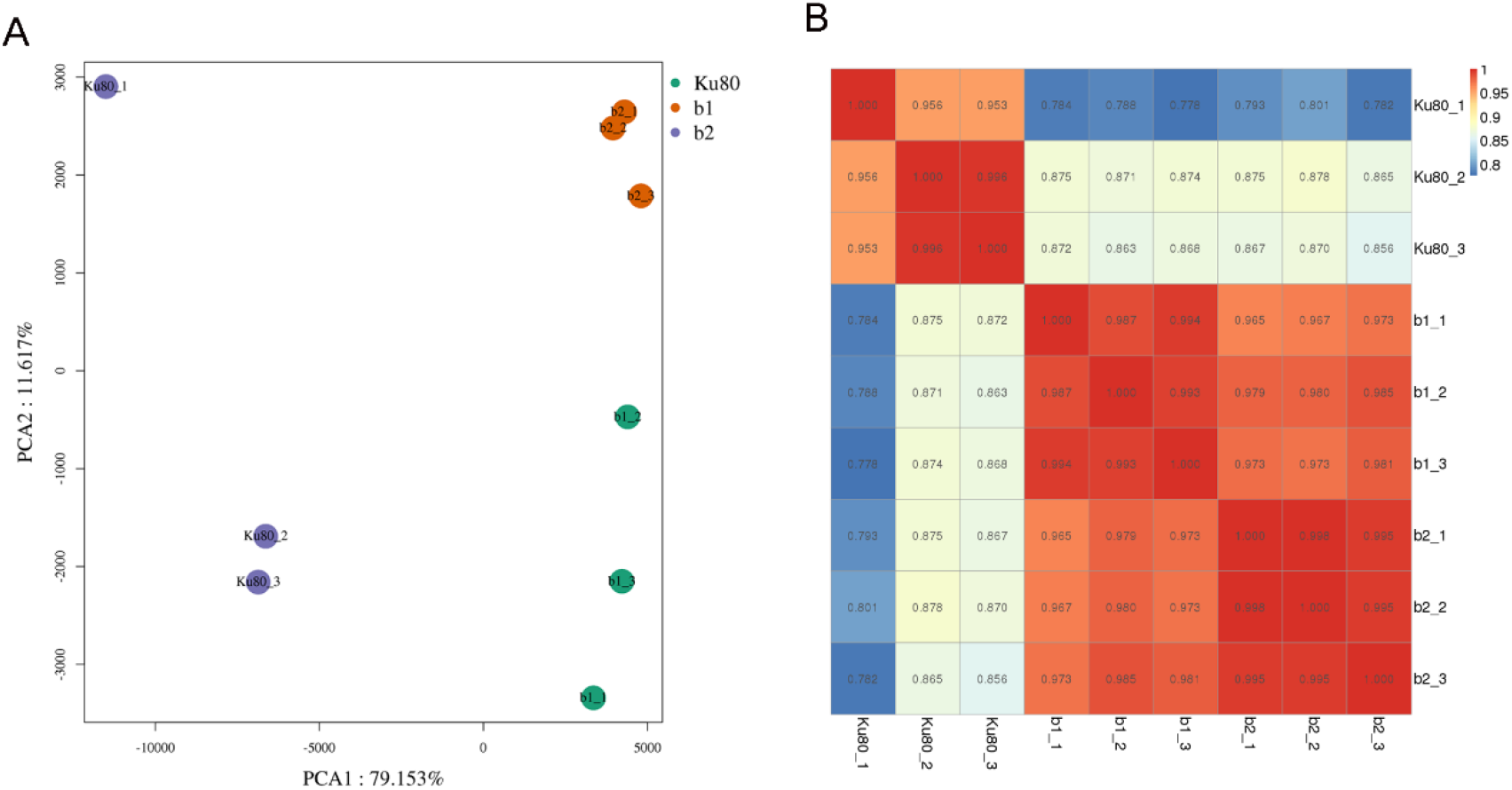
Principal component analysis (PCA) (A)and heatmap (B)showed the gene expression in background strains Ku80, mutant b1 and b2

**Fig. S3.**
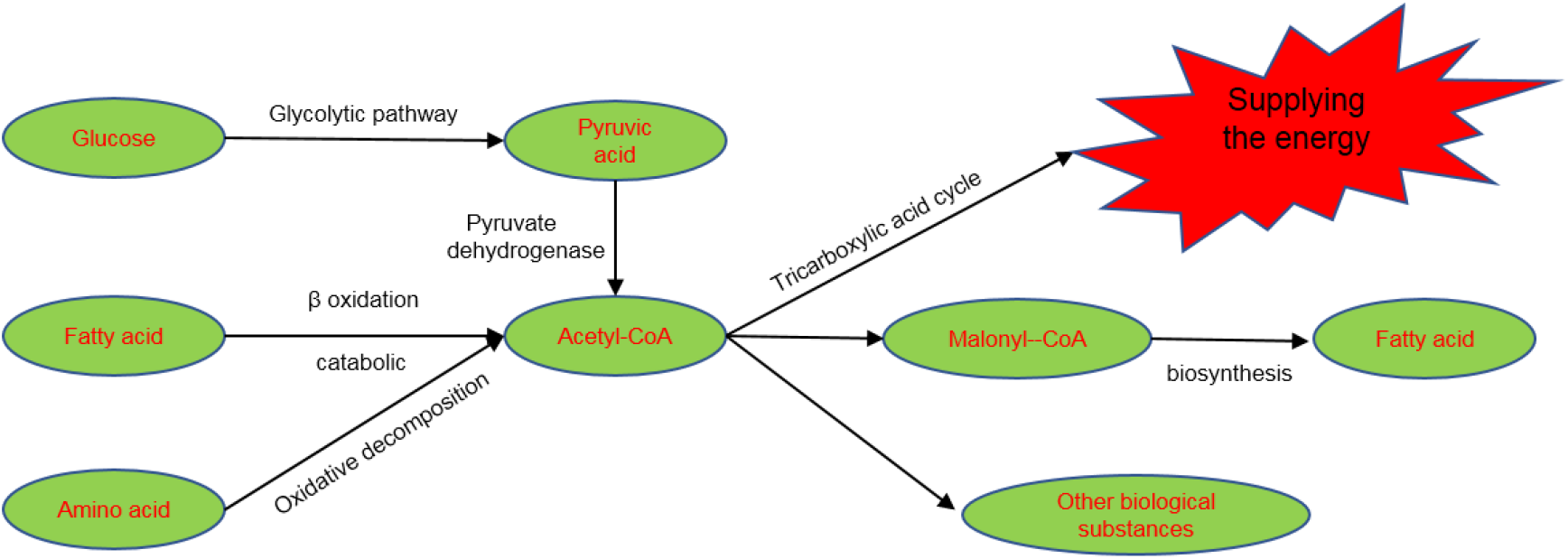
The metabolic pathway of acetyl CoA production and utilization in cells

**Fig. S4.**
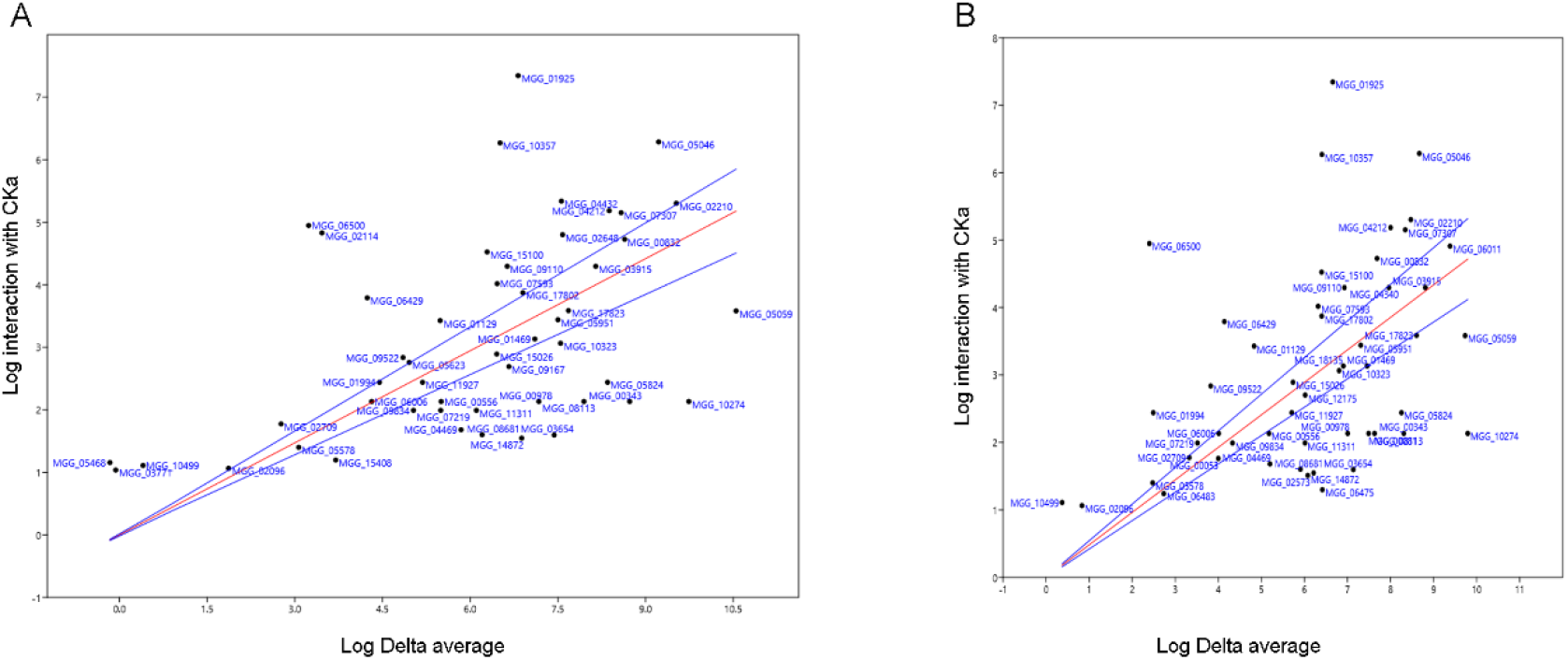
A and B same figures as in Figure 9 but with MGG gene codes

**Fig. S5.**
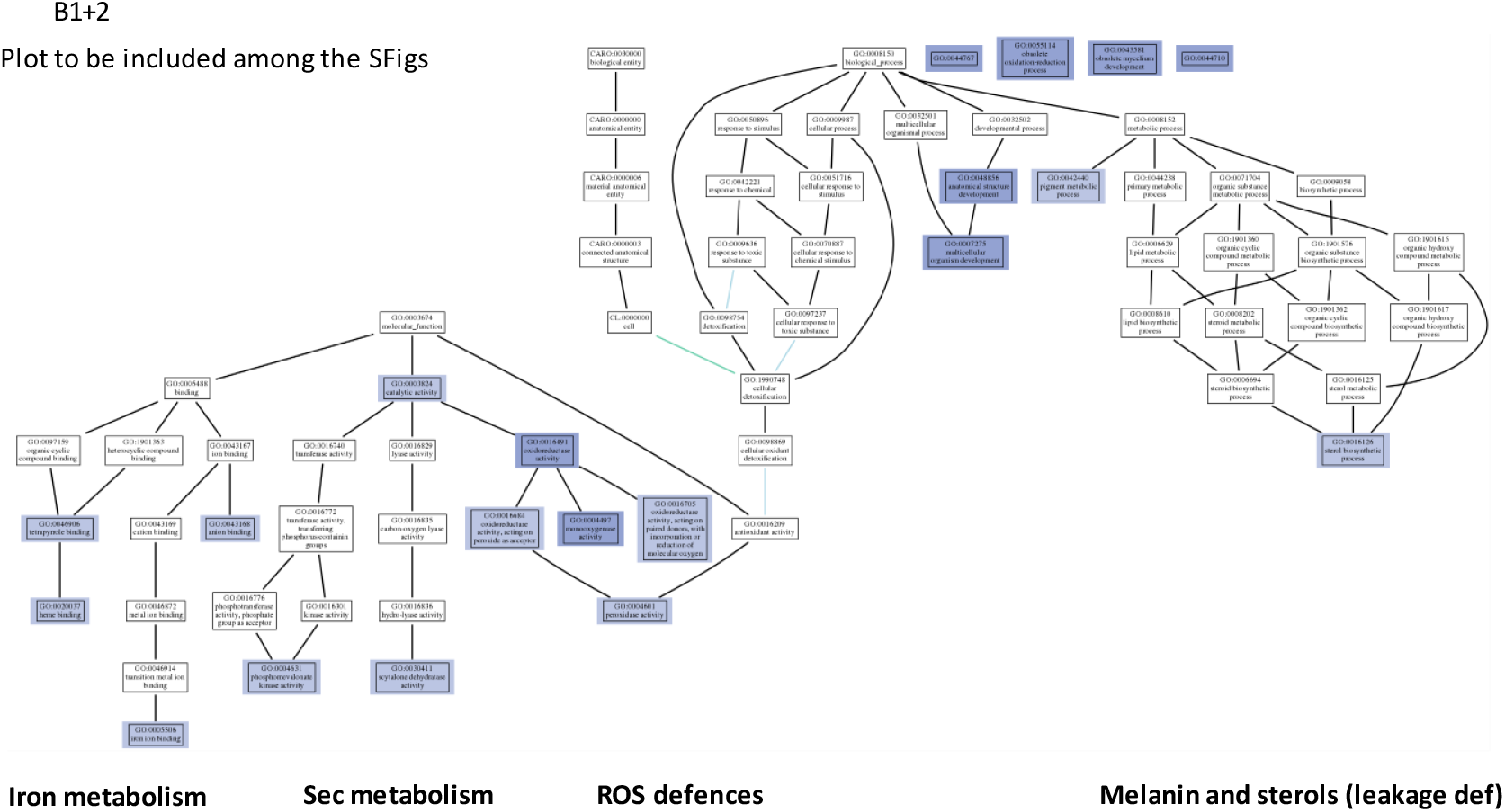
GO categories significantly enriched among the genes upregulated in both b1 and b2 mutants that are also significantly present in the CKa pulldown from a previous study

**Fig. S6.**
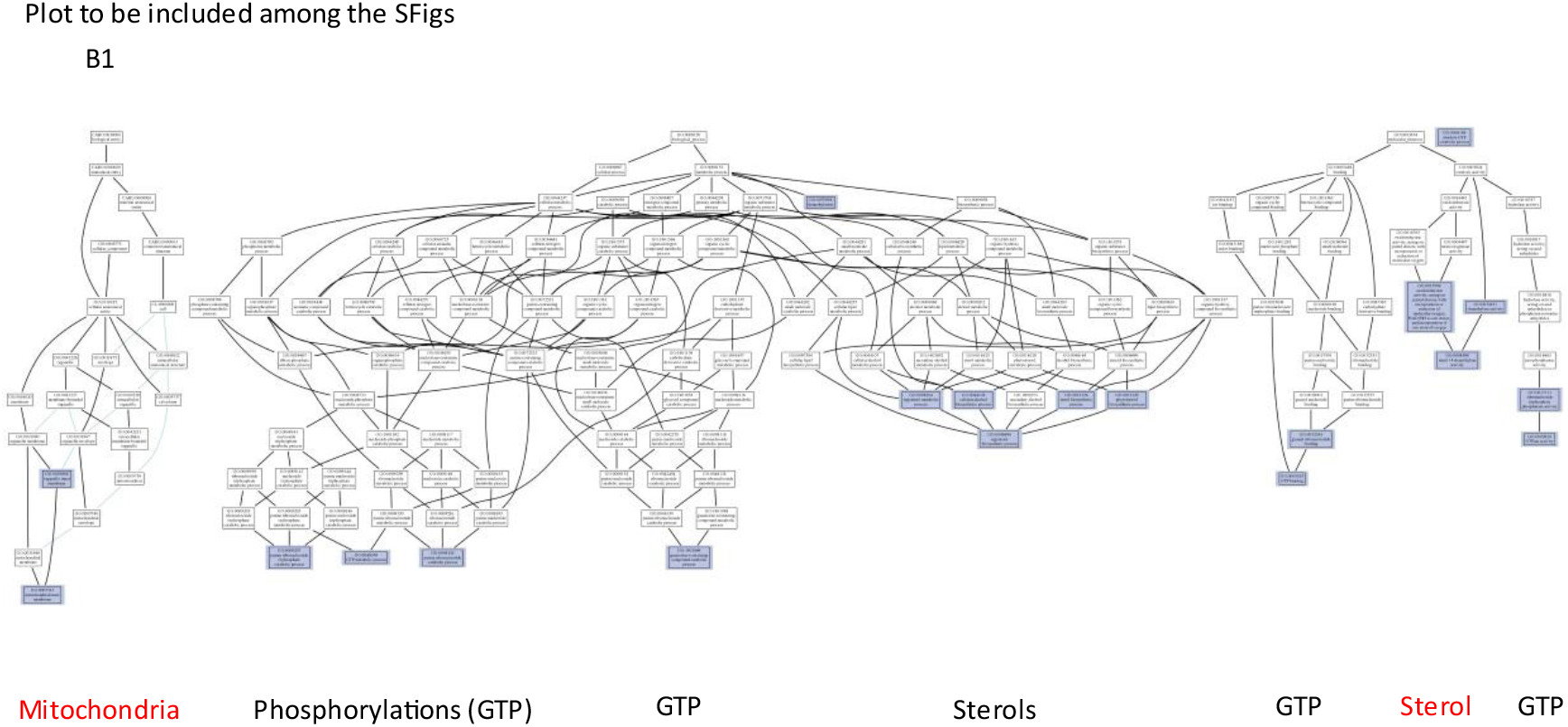
GO categories significantly enriched among the genes upregulated in the b1 mutant that are also significantly present in the CKa pulldown from a previous study. Mitochondria and sterol metabolisms are specially marked since many mitochondrial proteins were found in the CKa pulldown, and CK2 destabilization of alpha helixes of mitochondrial proteins could be essential for the efficient transport of these proteins into mitochondria. Sterol metabolism is also affected, which indicates that the b1 mutant is more negatively affected by membrane integrity.

**Fig. S7.**
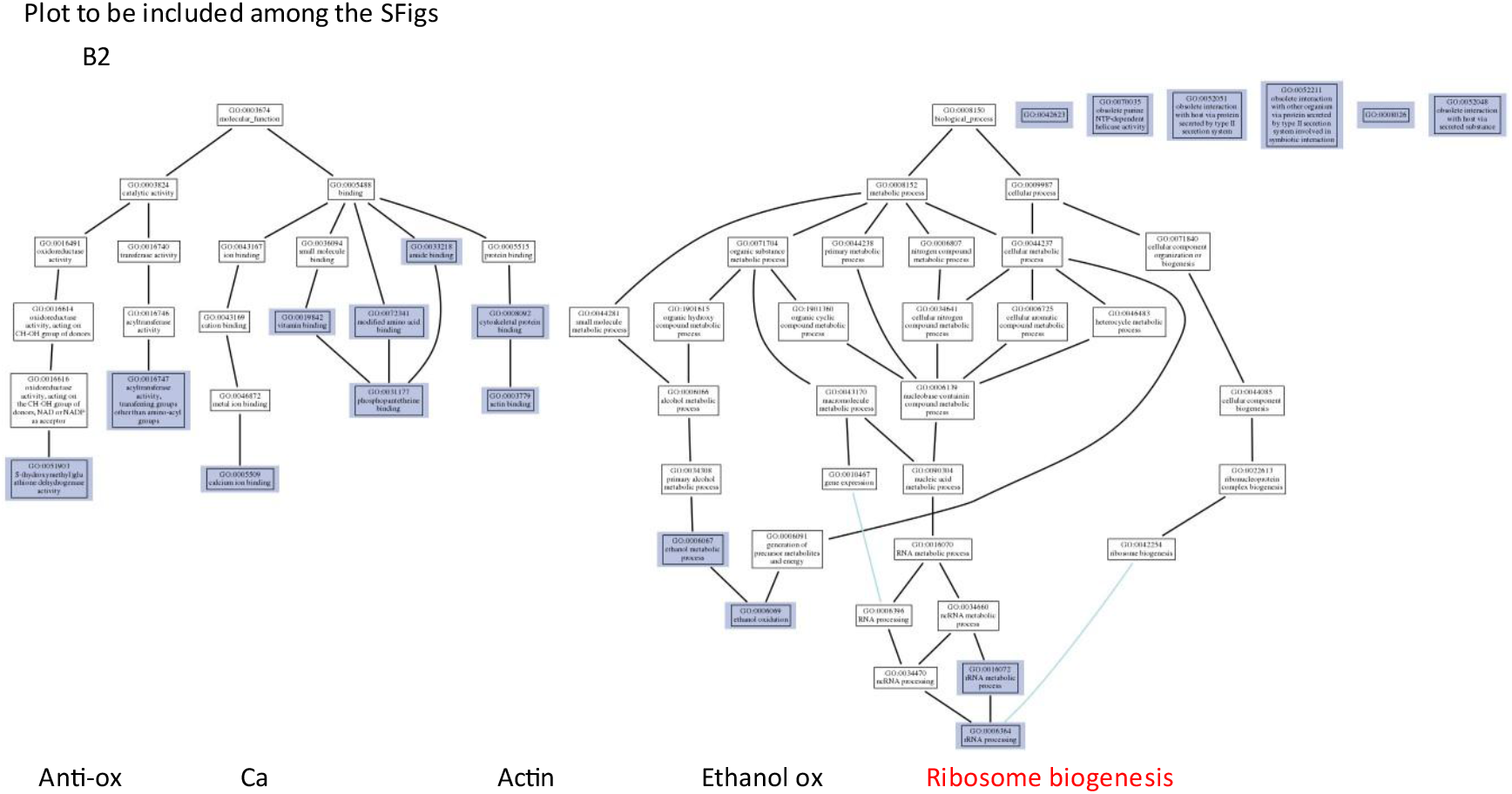
GO categories significantly enriched among the genes upregulated in the b2 mutant that are also significantly present in the CKa pulldown from a previous study. Ribosome biogenesis is especially marked since the CKa pulldown was especially enriched for intrinsically disordered ribosomal proteins indicating that the b2 mutant might be more affected in protein synthesis rate than the b1 mutant. That should be less important for responses to plants but could be very detrimental for fast fungal immunity responses(Ipcho et al., 2016).

**Fig. S8.**
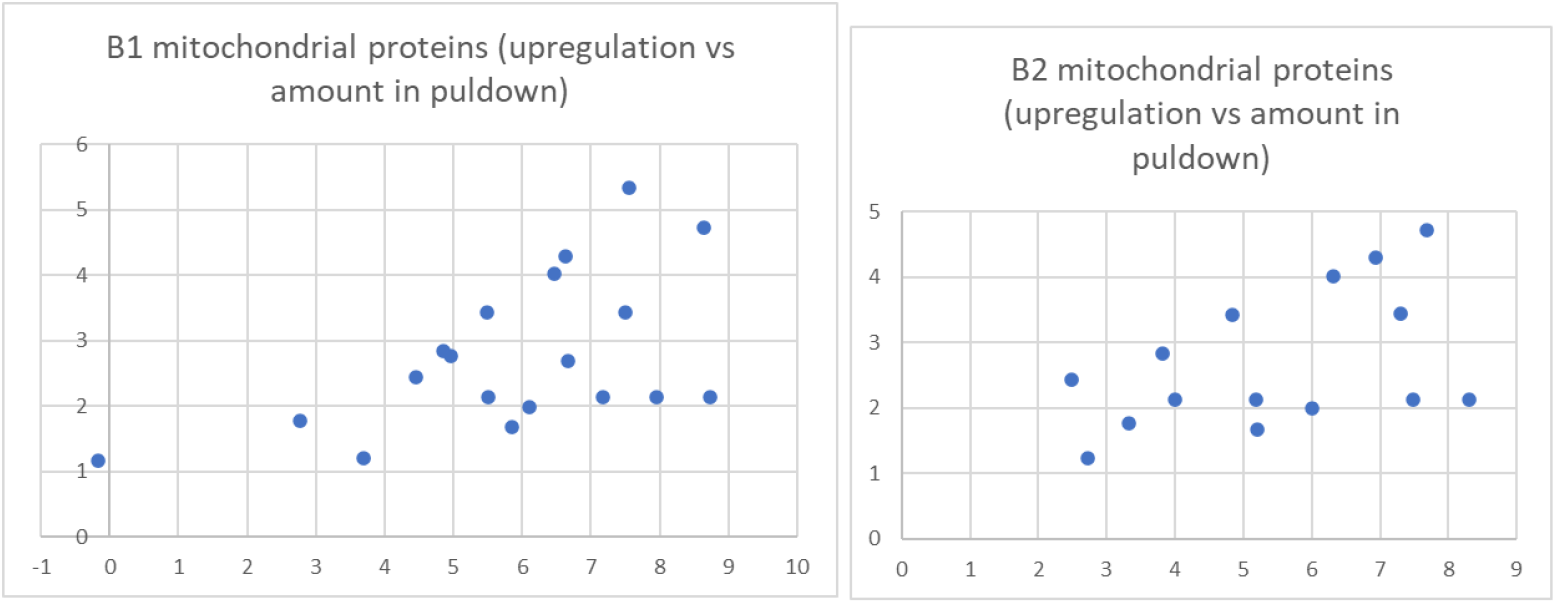
Absolute upregulation of mitochondrial genes versus the amount of the corresponding gene product in a CKa pulldown point towards a compensatory upregulation of most of these genes

**Table S1 information about transcriptome data**

Catalogue:

sheet1 Statistical table of clean data;

sheet2 Statistical table of comparison results;

sheet3 Distribution table of reads on gene functional elements;

sheet4 Location information table for new prediction genes;

sheet5 Location information table for annotated genes

sheet6 Quantitative analysis results of expression of all genes in different samples (FPKM value)

**Table S2 All different expression genes (DEGs) between b1, b2 mutants and background strains Ku80**

Catalogue:

sheet1 All different expression genes (DEGs) between b1 mutant and back ground strains Ku80; sheet2 the up-expression genes of DEGs in sheet 1;

sheet3 the down-expression genes of DEGs in sheet 1;

sheet4 All different expression genes (DEGs) between b2 mutant and back ground strains Ku80; sheet5 the up-expression genes of DEGs in sheet 4;

sheet6 the down-expression genes of DEGs in sheet 4

**Table S3 KEGG and GO analysis of DEGs in both b1 and b2 mutants**

Catalogue:

sheet1 KEGG enrichment of upregulated genes between b1 mutant and Ku80; sheet2 KEGG enrichment of down-regulated genes between b1 mutant and Ku80;

sheet3 KEGG enrichment of upregulated genes between b2 mutant and Ku80; sheet4 KEGG enrichment of down-regulated genes between b2 mutant and Ku80;

sheet5 GO annotation of upregulated genes between b1 mutant and Ku80;

sheet6 GO enrichment of upregulated genes between b1 mutant and Ku80; sheet7 GO annotation of down-regulated genes between b1 mutant and Ku80;

sheet8 GO enrichment of down-regulated genes between b1 mutant and Ku80; sheet9 GO annotation of upregulated genes between b2 mutant and Ku80;

sheet10 GO enrichment of upregulated genes between b2 mutant and Ku80; sheet11 GO annotation of down-regulated genes between b2 mutant and Ku80;

sheet12 GO enrichment of down-regulated genes between b2 mutant and Ku80;

**Table S4 different expression genes (DEGs) enriched in carbohydrate metabolism process in both deletion mutants**

Catalogue:

sheet1 GO enrichment of upregulated genes involving in carbohydrate metabolism in b1 mutant sheet2 GO enrichment of down-regulated genes involving in carbohydrate metabolism in b1 mutant sheet3 GO enrichment of upregulated genes involving in carbohydrate metabolism in b2 mutant sheet4 GO enrichment of down-regulated genes involving in carbohydrate metabolism in b2 mutant sheet5 upregulated genes involving in carbohydrate catabolic process in both mutants

sheet6 expression of upregulated genes involving in carbohydrate catabolic process in both mutants sheet7 down-regulated genes involving in carbohydrate catabolic process in both mutants

sheet8 expression of down-regulated genes involving in carbohydrate catabolic process in both mutants

**Table S5 different expression genes (DEGs) enriched in pyruvate metabolism in both deletion mutants**

sheet1 GO enrichment of upregulated genes involving in pyruvate metabolism in b1 mutant sheet2 GO enrichment of down-regulated genes involving in pyruvate metabolism in b1 mutant sheet3 GO enrichment of upregulated genes involving in pyruvate metabolism in b2 mutant sheet4 GO enrichment of down-regulated genes involving in pyruvate metabolism in b2 mutant sheet5 expression of pyruvate dehydrogenase and pyruvate transmembrane transporter in mutants

**Table S6 different expression genes (DEGs) enriched in fatty acid metabolism in both deletion mutants**

sheet1 GO enrichment of upregulated genes involving in fatty acid metabolism in b1 mutant sheet2 GO enrichment of down-regulated genes involving in fatty acid metabolism in b1 mutant sheet3 GO enrichment of upregulated genes involving in fatty acid metabolism in b2 mutant sheet4 GO enrichment of down-regulated genes involving in fatty acid metabolism in b2 mutant sheet5 expression of genes involving in fatty acid catabolic process in both mutants

sheet6 expression of genes involving in fatty acid fatty acid transmembrane transport in both mutants

**Table S7 different expression genes (DEGs) enriched in amino acid metabolism in both deletion mutants**

sheet1 GO enrichment of upregulated genes involving in amino acid metabolism in b1 mutant sheet2 GO enrichment of down-regulated genes involving in fa amino acid metabolism in b1 mutant sheet3 GO enrichment of upregulated genes involving in amino acid metabolism in b2 mutant sheet4 GO enrichment of down-regulated genes involving in amino acid metabolism in b2 mutant sheet5 genes enriched in cellular amino acid catabolic process (GO:0009063) in both mutants sheet6 genes enriched in aspartate family amino acid catabolic process (GO:0009068) in b1 mutant sheet7 genes enriched in serine family amino acid catabolic process (GO:0009071) in b1 mutant sheet8 genes enriched in glutamine family amino acid catabolic process (GO:0009065) in b1 mutant

**Table S8 different expression of transporter encoding genes in both deletion mutants**

sheet1 upregulated transporters in b1 mutant

sheet2 down-regulated transporters in b1 mutant

sheet3 upregulated transporters in b2 mutant sheet4 down-regulated transporters in b2 mutant

**Table S9 Significantly absolute upregulated genes for proteins found in a previous CKa pulldown that are predicted to be mitochondrial by DeepMito and their predicted mitochondrial location.**

**Table S10 Comparison of changes in regulation in CKb mutants with proteins found in a previous CKa pulldown also including a new prediction of mitochondrial proteins in the CKa pulldown using the web-tool DeepMito. Note all sheets with data in name are ordered in ID order so that the previous sheets can extract data dynamically from these using the VLOOKUP function in MSExcel.**sheet1 Up in b1 mutant D-ordsheet2 Up in b2 mutant D-ord sheet3 B1 plot

sheet4 B2 plot

sheet5 B1 B2 compared

sheet6 Mitochondria localization prediction table using DeepMito organized for paper

sheet7 CKa pulldown data from previous paper

sheet8 Disorder data from previous paper

sheet9 Mitochondria data from the DeepMito prediction of genes present in the CKa pulldown

## Author Contributions

RC and SO conceived and designed the experiments. LZ, CS, YZ, and WM performed the experiments. LZ, CS, RC, and SO analyzed the data and wrote the paper.

## Funding

This work was supported by the National Natural Science Foundation of China, Grant/Award Number: 32060597, and Jiangxi Provincial Natural Science Foundation, Grant/Award Number: 20212BAB215010.

## Notes

### Competing Interest Statement

The authors have declared no competing interest.

### Summary of Updates

no

## References

1. Skamnioti P, Gurr SJ. 2009. Against the grain: safeguarding rice from rice blast disease. Trends Biotechnol 27:141–50.

2. Wilson RA, Talbot NJ. 2009. Under pressure: investigating the biology of plant infection by Magnaporthe oryzae. Nat Rev Microbiol 7:185–95.

3. Miah G, Rafii MY, Ismail MR, Puteh AB, Rahim HA, Asfaliza R, Latif MA. 2012. Blast resistance in rice: a review of conventional breeding to molecular approaches. Molecular Biology Reports 40:2369–2388.

4. Bao J, Chen M, Zhong Z, Tang W, Lin L, Zhang X, Jiang H, Zhang D, Miao C, Tang H, Zhang J, Lu G, Ming R, Norvienyeku J, Wang B, Wang Z. 2017. PacBio Sequencing Reveals Transposable Elements as a Key Contributor to Genomic Plasticity and Virulence Variation in Magnaporthe oryzae. Mol Plant 10:1465–1468.

5. Wilson RA. 2021. Magnaporthe oryzae. Trends Microbiol 29:663–664.

6. Tucker SL, Talbot NJ. 2001. Surface attachment and pre-penetration stage development by plant pathogenic fungi. Annu Rev Phytopathol 39:385–417.

7. Foster AJ, Ryder LS, Kershaw MJ, Talbot NJ. 2017. The role of glycerol in the pathogenic lifestyle of the rice blast fungus Magnaporthe oryzae. Environ Microbiol 19:1008–1016.

8. Veneault-Fourrey C, Barooah M, Egan M, Wakley G, Talbot NJ. 2006. Autophagic fungal cell death is necessary for infection by the rice blast fungus. Science 312:580–3.

9. Saunders DG, Aves SJ, Talbot NJ. 2010. Cell cycle-mediated regulation of plant infection by the rice blast fungus. Plant Cell 22:497–507.

10. Xu JR, Staiger CJ, Hamer JE. 1998. Inactivation of the mitogen-activated protein kinase Mps1 from the rice blast fungus prevents penetration of host cells but allows activation of plant defense responses. Proc Natl Acad Sci U S A 95:12713–8.

11. Ebbole DJ. 2007. Magnaporthe as a model for understanding host-pathogen interactions. Annu Rev Phytopathol 45:437–56.

12. Adachi K, Hamer JE. 1998. Divergent cAMP signaling pathways regulate growth and pathogenesis in the rice blast fungus Magnaporthe grisea. Plant Cell 10:1361–74.

13. Xu JR, Hamer JE. 1996. MAP kinase and cAMP signaling regulate infection structure formation and pathogenic growth in the rice blast fungus Magnaporthe grisea. Genes Dev 10:2696–706.

14. Dagdas YF, Yoshino K, Dagdas G, Ryder LS, Bielska E, Steinberg G, Talbot NJ. 2012. Septin-mediated plant cell invasion by the rice blast fungus, Magnaporthe oryzae. Science 336:1590–5.

15. Zhang L, Zhang D, Chen Y, Ye W, Lin Q, Lu G, Ebbole DJ, Olsson S, Wang Z. 2019. Magnaporthe oryzae CK2 Accumulates in Nuclei, Nucleoli, at Septal Pores and Forms a Large Ring Structure in Appressoria, and Is Involved in Rice Blast Pathogenesis. Frontiers in Cellular and Infection Microbiology 9.

16. Zhang L, Zhang D, Liu D, Li Y, Li H, Xie Y, Wang Z, Hansen BO, Olsson S, Druzhinina IS. 2020. Conserved Eukaryotic Kinase CK2 Chaperone Intrinsically Disordered Protein Interactions. Applied and Environmental Microbiology 86.

17. Kumar V, Jain P, Venkadesan S, Karkute SG, Bhati J, Abdin MZ, Sevanthi AM, Mishra DC, Chaturvedi KK, Rai A, Sharma TR, Solanke AU. 2021. Understanding Rice-Magnaporthe Oryzae Interaction in Resistant and Susceptible Cultivars of Rice under Panicle Blast Infection Using a Time-Course Transcriptome Analysis. Genes 12:301.

18. El-Brolosy MA, Stainier DYR. 2017. Genetic compensation: A phenomenon in search of mechanisms. PLoS Genet 13:e1006780.

19. Kanehisa M, Furumichi M, Tanabe M, Sato Y, Morishima K. 2017. KEGG: new perspectives on genomes, pathways, diseases and drugs. Nucleic Acids Res 45:D353–d361.

20. Chen C, Chen H, Zhang Y, Thomas HR, Frank MH, He Y, Xia R. 2020. TBtools: An Integrative Toolkit Developed for Interactive Analyses of Big Biological Data. Mol Plant 13:1194–1202.

21. Shi L, Tu BP. 2015. Acetyl-CoA and the regulation of metabolism: mechanisms and consequences. Curr Opin Cell Biol 33:125–31.

22. Pronk JT, Yde Steensma H, Van Dijken JP. 1996. Pyruvate metabolism in Saccharomyces cerevisiae. Yeast 12:1607–33.

23. Poirier Y, Antonenkov VD, Glumoff T, Hiltunen JK. 2006. Peroxisomal beta-oxidation--a metabolic pathway with multiple functions. Biochim Biophys Acta 1763:1413–26.

24. Bhambra GK, Wang ZY, Soanes DM, Wakley GE, Talbot NJ. 2006. Peroxisomal carnitine acetyl transferase is required for elaboration of penetration hyphae during plant infection by Magnaporthe grisea. Mol Microbiol 61:46–60.

25. Aliyu SR, Lin L, Chen X, Abdul W, Lin Y, Otieno FJ, Shabbir A, Batool W, Zhang Y, Tang W, Wang Z, Norvienyeku J. 2019. Disruption of putative short-chain acyl-CoA dehydrogenases compromised free radical scavenging, conidiogenesis, and pathogenesis of Magnaporthe oryzae. Fungal Genet Biol 127:23–34.

26. Li Y, Zheng X, Zhu M, Chen M, Zhang S, He F, Chen X, Lv J, Pei M, Zhang Y, Zhang Y, Wang W, Zhang J, Wang M, Wang Z, Li G, Lu G. 2019. MoIVD-Mediated Leucine Catabolism Is Required for Vegetative Growth, Conidiation and Full Virulence of the Rice Blast Fungus Magnaporthe oryzae. Front Microbiol 10:444.

27. Ramos-Pamplona M, Naqvi NI. 2006. Host invasion during rice-blast disease requires carnitine-dependent transport of peroxisomal acetyl-CoA. Mol Microbiol 61:61–75.

28. Guerra B, Issinger OG. 2020. Role of Protein Kinase CK2 in Aberrant Lipid Metabolism in Cancer. Pharmaceuticals (Basel) 13.

29. Schwartz MP, Huang S, Matouschek A. 1999. The structure of precursor proteins during import into mitochondria. J Biol Chem 274:12759–64.

30. Savojardo C, Bruciaferri N, Tartari G, Martelli PL, Casadio R. 2020. DeepMito: accurate prediction of protein sub-mitochondrial localization using convolutional neural networks. Bioinformatics 36:56–64.

31. Kim D, Paggi JM, Park C, Bennett C, Salzberg SL. 2019. Graph-based genome alignment and genotyping with HISAT2 and HISAT-genotype. Nat Biotechnol 37:907–915.

32. Pertea M, Pertea GM, Antonescu CM, Chang TC, Mendell JT, Salzberg SL. 2015. StringTie enables improved reconstruction of a transcriptome from RNA-seq reads. Nat Biotechnol 33:290–5.

33. Pertea G, Pertea M. 2020. GFF Utilities: GffRead and GffCompare. F1000Res 9.

34. Liao Y, Smyth GK, Shi W. 2014. featureCounts: an efficient general purpose program for assigning sequence reads to genomic features. Bioinformatics 30:923–30.

35. Mortazavi A, Williams BA, McCue K, Schaeffer L, Wold B. 2008. Mapping and quantifying mammalian transcriptomes by RNA-Seq. Nat Methods 5:621–8.

36. Ben Salem K, Ben Abdelaziz A. 2021. Principal Component Analysis (PCA). Tunis Med 99:383–389.

37. Robinson MD, McCarthy DJ, Smyth GK. 2010. edgeR: a Bioconductor package for differential expression analysis of digital gene expression data. Bioinformatics 26:139–40.

38. Ipcho S, Sundelin T, Erbs G, Kistler HC, Newman MA, Olsson S. 2016. Fungal Innate Immunity Induced by Bacterial Microbe-Associated Molecular Patterns (MAMPs). G3 (Bethesda) 6:1585–95.

